# Self-generation of goal-directed choices in a distributed dopaminergic and prefrontal circuit

**DOI:** 10.1101/2022.05.19.492598

**Authors:** E Bousseyrol, S Didienne, S Takillah, C Solié, M Come, Ahmed Yahia T, S Mondoloni, E Vicq, L Tricoire, A Mourot, J Naudé, Ph Faure

**Affiliations:** Sorbonne Université, INSERM, CNRS, Neuroscience Paris Seine - Institut de Biologie Paris Seine (NPS - IBPS), 75005 Paris, France.; Brain Plasticity Unit, CNRS, ESPCI Paris, PSL Research University, 75005 Paris, France; CNRS, Université de Montpellier, INSERM - Institut de Génomique Fonctionnelle, 34094 Montpellier, France.

## Abstract

Goal-directed choices that are not triggered by external cues arise from internal representations of the outcomes. The use of a stimulus to specify when to act, which option to take, or whether to explore, has led to consider the reward circuit as a feedforward set of modules carrying independent computations. Here, we develop an uncued task in which mice self-determine the initiation, direction, vigor and pace of their actions based on their knowledge of the outcomes. Using electrophysiological recordings, pharmacology and optogenetics, we identify a sequence of oscillations and firing in the ventral tegmental area (VTA), orbitofrontal (OFC) and prefrontal cortices (PFC) that co-encodes and co-determines self-initiation and choices. This sequence appeared with learning as an unguided realignment of spontaneous dynamics. The interactions between the structures depended on the reward context, in particular regarding the uncertainty associated with the different options. We suggest that self-generated choices arise from a distributed circuit based on an OFC-VTA core setting whether to wait or to initiate actions, while the PFC is specifically engaged by reward uncertainty to participate in both the selection and pace of actions.

**Highlights:**

- Self-paced actions arise from contextual reorganization of mesocortical dynamics.
- VTA, PFC and OFC complementarily encode predictions and errors about outcomes.
- Distributed firing-then-oscillations dynamics set the goal, initiation and pace of actions.
- VTA and PFC antagonistically promote and inhibit motivation by reward uncertainty.

## Introduction

A major distinction is made between respondent and operant behaviors ^1, 2^. Animal behavior is said to be respondent when actions are triggered by an external stimulus. Nevertheless, animals often base their decisions solely on internal representations of their goals. In such case, the behavior is said to be operant but also “self-paced”, “self-generated” or “self-initiated” ^1, 3^. This includes decisions such as when to act, but also such as which option to take in cases where no stimulus indicates each option’s value. Decision-making in the absence of cues thus necessitates to compare internal memories of the values associated with different options. In particular, when faced with several alternatives, animals do not always exploit the option associated with the highest expectation of reward, but instead explore less rewarded options, in order to gain information ^4^. Indeed, while exploration might be guided by novel or salient cues, it can also be a consequence of an internal exploratory mode ^4^. However, most of our knowledge on decision-making has been obtained with respondent, stimulus-triggered behaviors that are most reproducible and amenable to statistical analyses ^1, 2^. Hence, the neural mechanisms by which animals initiate goal-directed actions, decide between them, and explore potentially informative ones in the absence of external cues, is far from being understood.

The choice of the experimental paradigm through which the neural mechanisms of goal-directed choice are probed is not neutral. In respondent behavior, the stimulus constitutes an identified input to the nervous system and the action its output ^5^. In this framework, it is tempting to treat the decision circuit as a feedforward process. Hence, a long-standing question has consisted in assigning a modular and sequential role to each stage of the circuit ^6^, e.g. to the ventral tegmental area (VTA) midbrain dopamine nucleus, the frontal areas (i.e. prefrontal and orbitofrontal cortices) and basal ganglia. This is illustrated by the reinforcement-learning computational theory, which identifies phasic release of dopamine (DA) with a reward prediction error (RPE), i.e. the comparison between actual and expected reward. DA RPE would constitute a teaching signal for learning the appropriate stimulus-action responses ^7, 8^. In this theory, DA would only play an indirect role in behavior, affecting subsequent trials of a task rather than the current one. In the modular view, cortices provide subcortical areas with information about the current state and options of the environment and the basal ganglia then select among these options to initiate the goal-directed action ^3, 9^. Hence, frontal cortices would, like DA, only indirectly affect decisions, by signaling sensory states, beliefs and possibilities to the basal ganglia, and/or by conveying value predictions to be compared with actual outcomes by DA neurons ^10, 11^.

However, both dopamine and frontal areas have been described as having more direct roles in decision-making, and the basal ganglia might not be the only locus of action selection ^12^. Good-based models place the choice process at the level of the OFC/PFC ^13^, as frontal cortices not only represent the value of potential options but also actively compare between them ^6, 14^. It is however unclear how economic choice is independent from action planning ^14^ or result from recurrent computations incorporating motor parameters of action execution back into a recurrent decision process ^15^. Theories on cognitive control assign a top-down, potentially inhibitory role to the OFC and PFC through computing goals, plans and task rules, i.e. higher-order decision-making ^16^. These different accounts all point at a direct, active role of frontal areas on ongoing choices. Phasic DA also exerts an impact on the ongoing behavior and on self-paced actions, by modulating the vigor of actions leading to reward ^17, 18^. Mixed results have been obtained on DA facilitating action initiation itself, depending on the type of task, DA nuclei, and intensity of DA manipulation ^19–22^. Given the respective roles of DA and frontal cortex in value-based decision making, selection and action initiation, the question of the coordination between nodes of this meso-cortical circuit arises, in learning processes, in decision-making and exploration, and in motor execution.

Overall, studies focusing on prefrontal modulation of the VTA have focused on learning ^10, 11, 23, 24^ while work assessing dopamine modulation of frontal cortices have pointed at decision-making ^25–27^. As the mesocortical circuit is a loop rather than a feedforward process ^6, 28^, an integrated view is thus lacking. Concurrent recordings of VTA and frontal cortices ^29, 30^ have shown synchrony between these structures but have not delved into causal mechanisms. More importantly, these studies have focused on the role of DA or cortical dynamics in cue-guided behaviors ^10, 11, 23–25, 29, 30^, or, when assessing goal-directed actions, did not examine the respective roles of these structures in action self-initiation ^26, 27^. In the present study, we used an experimental paradigm in which mice must perform a sequence of choices to obtain rewards associated with intracranial self-stimulation ^31–33^. This protocol satisfies a series of requirements that have proven difficult to address altogether: simultaneously monitoring action selection, action execution, and electrophysiological activity of the mesocortical circuit, in an environment with controlled rewards and exploration. Most importantly, mice generate a vast number of “template” bouts of locomotion in this task ^32^, at a constant motivational level insured by intracranial self-stimulation.

In this task, we show that goal-directed actions from one rewarded location to the next consisted in stereotyped, ballistic (bell-shaped speed profiles) movements that nonetheless presented characteristic variabilities in their timings or directions. Mouse behavior displayed hallmarks of self-generated actions, with initiation, pace and decisions underdetermined by environmental information, but influenced by internal representations of the reward context. We further show that the VTA, PFC and OFC coordinated their activities into a sequence of distributed firing and oscillations, associated with the goal-directed actions. Such sequence emerged with learning as a reorganization of existing dynamics locking on behaviorally-relevant timings, without being triggered by a stimulus. Combining electrophysiology with optogenetic and pharmacological manipulations, we unveil that OFC-VTA interactions set self-initiation in each reward context, with a more specific involvement of the PFC during decisions under uncertainty and exploration. We also suggest that the VTA and PFC can act in synergy to self-pace the actions, but may have antagonistic roles in pondering the influence of uncertainty on choices, in particular for exploration. Our study highlights how the mesocortical circuit self-generates decisions through distributed yet distinct computations, by reorganizing itself in a context-dependent manner.

## Results

### Mouse actions underdetermined by environmental cues, but shaped by internal representations, indicate self-paced decisions

To decipher the neural bases of self-paced decisions, we used a spatial version of a multi-armed bandit task adapted to mice ^31–33^. Three equidistant locations explicitly marked in an open-field were associated with rewards delivered as intra-cranial self-stimulations (ICSS) in the medial forebrain bundle (MFB) ^34^ (Figure 1A left). The MFB ICSS eliminates the need for food restriction and the associated satiation level, which is known to affect decisions, in particular under reward uncertainty ^35^. Mice could not receive two consecutive ICSS at the same location, they therefore alternated between rewarding locations (Figure 1A right). Mice were initially trained in a deterministic context (D) in which all locations were associated with a certain ICSS delivery (P=100%). Then, mice underwent a probabilistic (P) context, in which each location was associated with a different probability of ICSS delivery (here; P=100%, 50% and 25%; Figure 1A), to assess how self-paced actions and meso-cortical representations depend on outcome expectations. Their behavior in the task after learning in the deterministic context thus consisted in sequences of trials in-between rewarding locations. First, on each rewarded location, mice could either circle forward along the three locations, or perform U-turns by coming backwards, at trial i+1, to the i-1 location (Figure 1B). Second, a trial was characterized by a “template” bout of locomotion: a movement initiation towards the next location, an acceleration followed by deceleration and a pause at the next location (Figure 1C left). Execution of this ballistic (bell-shaped) velocity profile was characterized by an important trial-to-trial variability, notably regarding the time during which animals dwelled at a rewarded location before initiating a new trial, the time to reach maximal speed, and the overall time to goal (from one location entry to the next, Figure 1C right). As these successive timings were nested within the behavioral sequence, their variability could correspond to a global decision on the overall timing of the trial. In such case, variability of intermediate timings would be correlated, either positively (with the first timings already explaining the subsequent variability) or negatively (with the variability in each timing compensating in a zero-sum fashion). Alternatively, a successive, independent addition of variability at each stage of the trial could sign distinct decisions about movement direction, initiation and vigor. In the deterministic context, the direction (U-turn or Forward) impacted both on the dwell duration - an early trial timing, and finally on the time to goal (Figure 1D). Multiple linear regressions (with orthogonalized predictors, model p-value <0.05 for every animal, see Methods) showed that the trajectory direction, the dwell time and the time to maximal speed all had (independently from each other) significant impact on the time to goal (Figure 1E). Hence, each stage of a trial (trajectory direction, initiation, and vigor) presented a successive addition of independent variability suggesting that trajectory direction, initiation, and vigor all constituted decisions for the animals ^1, 3^. Self-paced decisions depend on the representation of the potential outcome. We thus assessed whether the stages of the trial were affected by the animals’ internal representations of the potential rewards, by comparing behaviors in two reward contexts, the deterministic context and the probabilistic context (with different probabilities of ICSS delivery at each location, P=100%, 50% and 25%; Figure 1A). After training in the probabilistic context, the time to goal and dwell time increased compared to the deterministic context, even when considering only forward trajectories, suggesting a decreased motivation due to a decrease in reward delivery frequency in the probabilistic context (Figure 1F). The proportion of U-turns increased as well, reflecting an adjustment of the trajectory directions to the respective payoffs of the locations: while mice visited the three locations uniformly in the deterministic context, in the probabilistic context they visited more often the locations associated with the highest ICSS probabilities (i.e. p_100_ and p_50_, Figure 1G). Indeed, choices were modulated by reward locations in the probabilistic context but not in the deterministic context, and locations differentially affected choices in the deterministic and probabilistic contexts. Hence, mouse behavior displayed hallmarks of self-paced decisions: the direction, initiation and vigor of actions presented independent variability, were not determined by a stimulus cue, but rather were influenced by the potential outcomes of the actions.

**Figure 1:**
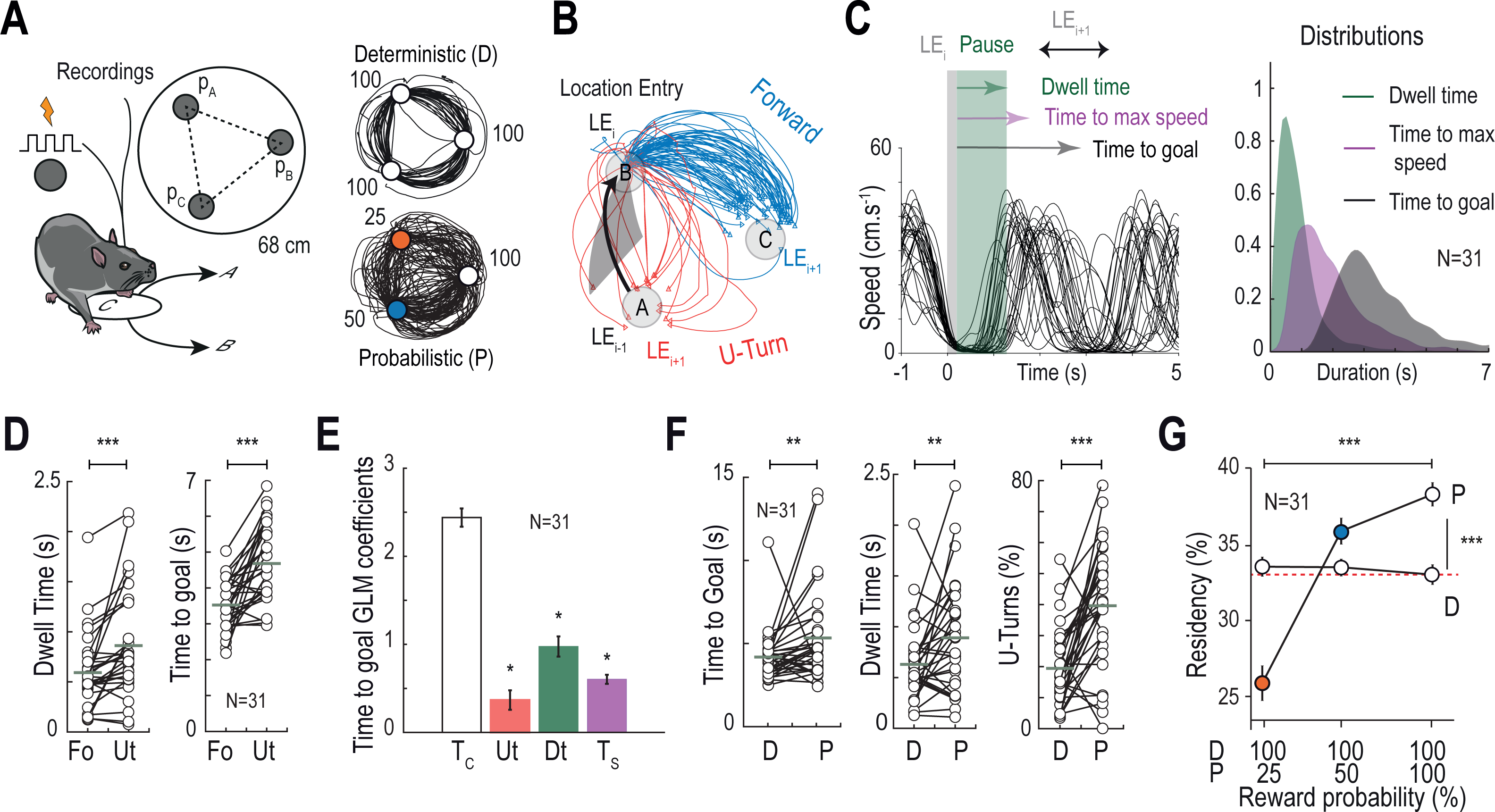
Self-paced decisions in a mouse spatial task based on intracranial self-stimulations. **(A)** Left: illustration of the task design. Three explicit square locations were placed in the open field (0.68-m diameter), forming an equilateral triangle (35-cm side). Each location is associated with a given probability of intracranial self-stimulation (ICSS) delivery (P=100%, 50%, 25%) when the mouse is detected in the location area. Animals could not receive two consecutive ICSS at the same location. Right: examples of trajectories (5 min) showing that mice alternated between rewarding locations in the deterministic (D) and probabilistic (P) contexts. **(B)** Animals varied between forward trajectories (Fo), in which mice keep the direction of their last choice, thus performing ‘A-B-C’ sequences; and U-turn trajectories (Ut), in which mice went backward after their previous location, corresponding to A-B-A sequences. **(C)** Left: examples of instantaneous speed of one mice after learning in the D setting, showing that the animals almost stopped at the time of the reward after location entry (henceforth called ”location entry” LE), stayed immobile for a short dwelling period, then accelerated toward their next location. These bouts of activity can be described using three observables, the dwell time (time from ICSS delivery during which speed is less than 10 cm/s), the time to the maximal speed, and the total time to goal from one location to the next. Right: distribution for these three parameters, for all trials of all mice, at the end of the D setting. **(D)** In the D context, dwell time and time to goal were higher for mice performing U-turn (Ut) trajectories compared to Forward (Fo) trajectories. (Dwell time: Δ=0.27s, paired two-sided Wilcoxon signed rank test, W_(30)_=-38, p<0.001; Time to goal: Δ=1.16s, paired Student t-test, T_(30)_=-4.95, p<0.001). Grey horizontal bars represent the means. **(E)** Coefficients of the generalized linear model of time to goal in the D context: constant term (Tc), U-turn (categorical variable, either U-turn or forward) (Student t-test T_(30)_=3.31, p=0.0024), dwell time (Dt) (two-sided Wilcoxon signed rank test W_(30)_=496, p<0.001) and time to maximal speed (Ts) (Student t-test T_(30)_=11.76, p<0.001). Vertical bars represent SEM. Stars (*) indicate a significant impact on the time to goal. **(F)** Time to goal, dwell time, and proportion of U-turns increased in the probabilistic (P) context compared to the deterministic (D) context. (Time to goal: paired two-sided Wilcoxon signed rank test, Δ=1.14s, W_(30)_=104, p=0.005; Dwell time for all trajectories: paired Student t-test T_(30)_=-2.64, p=0.01, Δ=0.26s; Dwell time for forward trajectories only: Student t-test T_(30)_=-3.27, p=0.0027, for time to goal; U-Turn: paired Student t-test, Δ=0.20s, T_(30)_=-4.64, p<0.001) Grey horizontal bars represent the means. **(G)** Proportion of choices of the three rewarding locations, as a function of reward probability in the P and D contexts. Effect of the P context on choice distribution: F_(30,2)_=48.5, p<0.001; Same for D context: F_(30,2)_=0.2, p=0.81, One-way ANOVAs; Effect of probabilities on choices in the P context: F_(1,2)_=31.8, p<0.001, two-way ANOVA. Vertical bars represent SEM.

A distributed mesocortical sequence associated with self-generated decisions depends on reward context As self-generated behaviors are caused by changes in the internal state of the animals, rather than by external changes in the environment ^1, 3^, we next characterized the mesocortical dynamics associated with self-paced decisions. We recorded from putative dopamine neurons (pDAn, n=136 neurons) in the VTA from WT mice (n=12 mice) using extracellular multi-electrodes (Figure 2A Left). All neurons met the electrophysiological and pharmacological criteria used to identify dopamine cells in vivo ^36, 37^ (Supp. Figure 1, Methods). In another group of mice, bipolar electrodes were also chronically implanted bilaterally into the prefrontal (PFC) and orbitofrontal (OFC) cortices (n=23 mice), allowing to record both extracellular field potentials (EFP) and population spiking activity (Figure 2A right, Supp. Figure 1, Methods).

**Figure 2:**
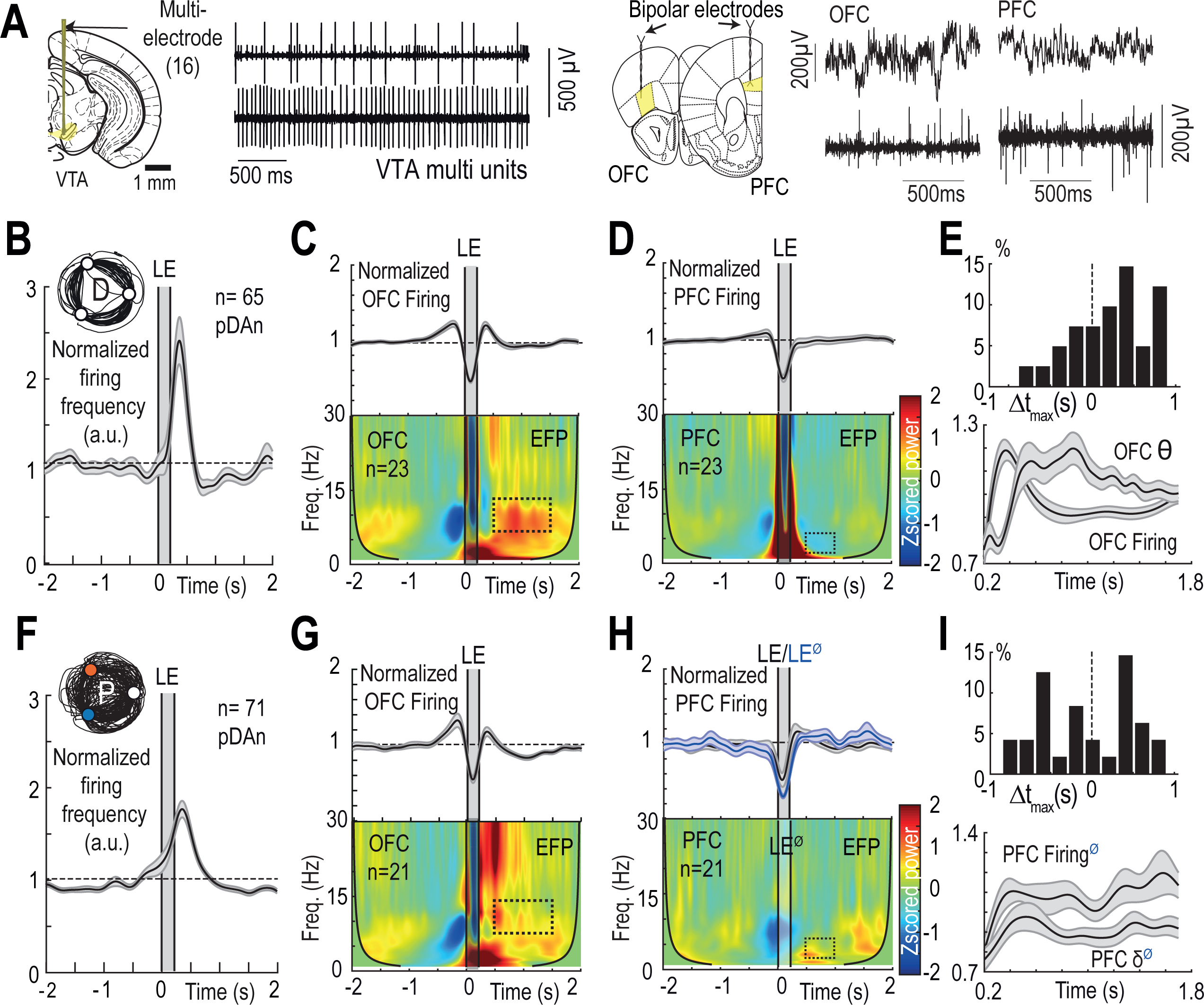
VTA, PFC and OFC during deterministic and probabilistic reward contexts. **(A)** Left: filtered (600-6000 Hz) extracellular recordings in the VTA showing multiple unit activity. Right: Extracellular recordings in the OFC and PFC showing extracellular field potential (EFP, filtered 0.1-300 Hz) (top) or population activity (filtered 600-6000 Hz) (bottom). **(B)** Normalized firing frequency from VTA pDAn around the time at which the animal is first detected in the location (LE), i.e. reward delivery in the D context. (pDAn mean firing over 0.3-0.8s, two-sided Wilcoxon-Mann-Whitney test W_(64)_=1572, p=0.001). Data are presented as mean ± SEM. **(C)** Top: Normalized firing frequency from OFC population (mean ± SEM) around the time at which the animal is first detected in the location in the D context (i.e. reward delivery). (OFC mean firing over 0.3-0.8s, Student t-test T_(39)_=2.57, p=0.01). Bottom: 0-30Hz range time-resolved power spectral density (PSD, using a complex Morlet wavelet transform) of OFC EFP around reward delivery. PSD is Zscored over the 2s period preceding the location entry (LE). (Mean OFC θ 7-14Hz power over 0.5-1.5s power Student t-test T_(22)_=5.12, p<0.001). **(D)** Same as (C) for the PFC. **(E)** Top: Time lag between the maximal OFC θ oscillation power and the maximal OFC firing. (Time of the maximum θ power minus time of maximum firing: two-sided Wilcoxon-Mann-Whitney test U_(78)_=1371, p=0.002). Bottom: Superposition of OFC θ oscillation and population firing frequency. Data are presented as mean ± SEM. **(F,G,H)** same as (B,C,D) but in the P context. (F) Mean pDAn firing frequency over 0.3-0.8s: Student t-test T_(74)_=7.61, p<0.001. Difference compared to the D setting shown in (B): ns two-sided Wilcoxon-Mann-Whitney test U_(64)_=4345, p=0.32. (G) Difference between P and D settings: (Top) OFC mean firing frequency during 0.3-0.8s: ns Student t-test T_(91)_=0.56, p=0.6. (Bottom) OFC mean θ power during 0.5-1.5s: ns Student t-test T_(42)_=1.12, p=0.3. (H) (Top) PFC mean firing over 0.5-1s post-omission: two-sided Wilcoxon signed rank test W_(49)_=866, p=0.03. (Bottom) The PFC PSD is centered on reward omissions (LE^Ø^) to prevent the ICSS artefact from obscuring the low frequency power. PFC δ mean power during 0.5-1s window post-omission: Student t-test T_(20)_=2.07 p=0.05; difference with the D context: two-sided Wilcoxon-Mann-Whitney test U_(20)_=355, p<0.001. **(I)** (Top) Time lag between the maximal PFC δ oscillation power and the maximal PFC firing, post-omission only. (Time of the maximum δ power minus time of maximum firing: ns paired two-sided Wilcoxon signed rank test W_(47)_=2121). Bottom: Superposition of PFC δ oscillation power and population firing frequency.

At the end of the deterministic context, pDAn (n=65 cells) emitted bursts of action potentials early in the trials (henceforth called “early bursting”), after the location entry and ICSS delivery, during the dwelling period (Figure 2B). Early bursting in VTA pDAn was concurrent with an early increase in OFC population firing (Figure 2C top). After the dwell time, when the animals started to move towards the next location (0.5-1.5s window), oscillation power in the θ band (7-14 Hz) increased in the OFC EFP (Figure 2C bottom). In contrast, no specific activity was observed in the PFC around the location entry, neither in the population spiking nor in the EFP (Figure 2D). Consistent with this temporal order, OFC firing and θ oscillations in the OFC displayed a lagged cross-correlation (Figure 2E below), indicating a transition from increased spiking to θ oscillations around the time of self-initiation of the trial by the mice. Although some of the oscillations in the PFC and OFC co-occurred, we did not observe any increase in coherence, in none of the contexts (deterministic or probabilistic, Supp. Figure 2).

We next asked how mesocortical activity changed with the reorganization of animals’ choices when faced with the uncertainty of probabilistic reward delivery. pDAn early bursting was still present at the end of the probabilistic context and the increase in firing frequency (Figure 2F) was not different from what was observed in the deterministic context. The parallel increase in OFC firing during dwelling, and latter power increase in θ oscillations, were similar in both contexts (Figure 2G). By contrast, in the probabilistic context, an activity consisting in an increase in both population firing (0.5-1s window after location entry, post-omission only) and δ (3-6 Hz) oscillation power (0.5-1s window after location entry, post-omission only emerged in the PFC (Figure 2H). No preferential temporal lag was observed between PFC firing and δ oscillations after the location entry (Figure 2I), suggesting an absence of a clear temporal order between firing and oscillations. Overall, the mesocortical network, classically associated with stimulus-triggered decision-making ^10, 23, 24^, was also recruited during self-generated decisions, but with an involvement of the PFC specifically during decisions under uncertainty. Our results are consistent with the view that self-generated decisions do not need a separate neural substrate but might reuse the neuronal circuits and computations of cue-guided behavior, yet in an ‘internally generated’ mode of brain dynamics^38^, with the PFC exerting a context-dependent influence when choice values are uncertain ^39^.

### A self-generated mesocortical sequence emerges with learning as a reorganization of existing dynamics

The influence of reward context on VTA, PFC and OFC activities suggests that the mesocortical dynamics associated with self-paced actions reorganizes based on the actions’ outcomes. We thus investigated the emergence of this dynamics with learning in naïve mice within the deterministic context. The total number of reward locations visited per session increased throughout the sessions (Figure 3A right), confirming place reinforcement. The averaged time to goal (Figure 3B left) and the dwell time (Figure 3B middle left) decreased accordingly, while the maximal speed increased (Figure 3B middle right). Finally, the proportion of U-turn decreased (Figure 3B right), indicating that mice learned to optimize their trajectories, and reduced the motor cost associated with U-turns ^31^.

**Figure 3:**
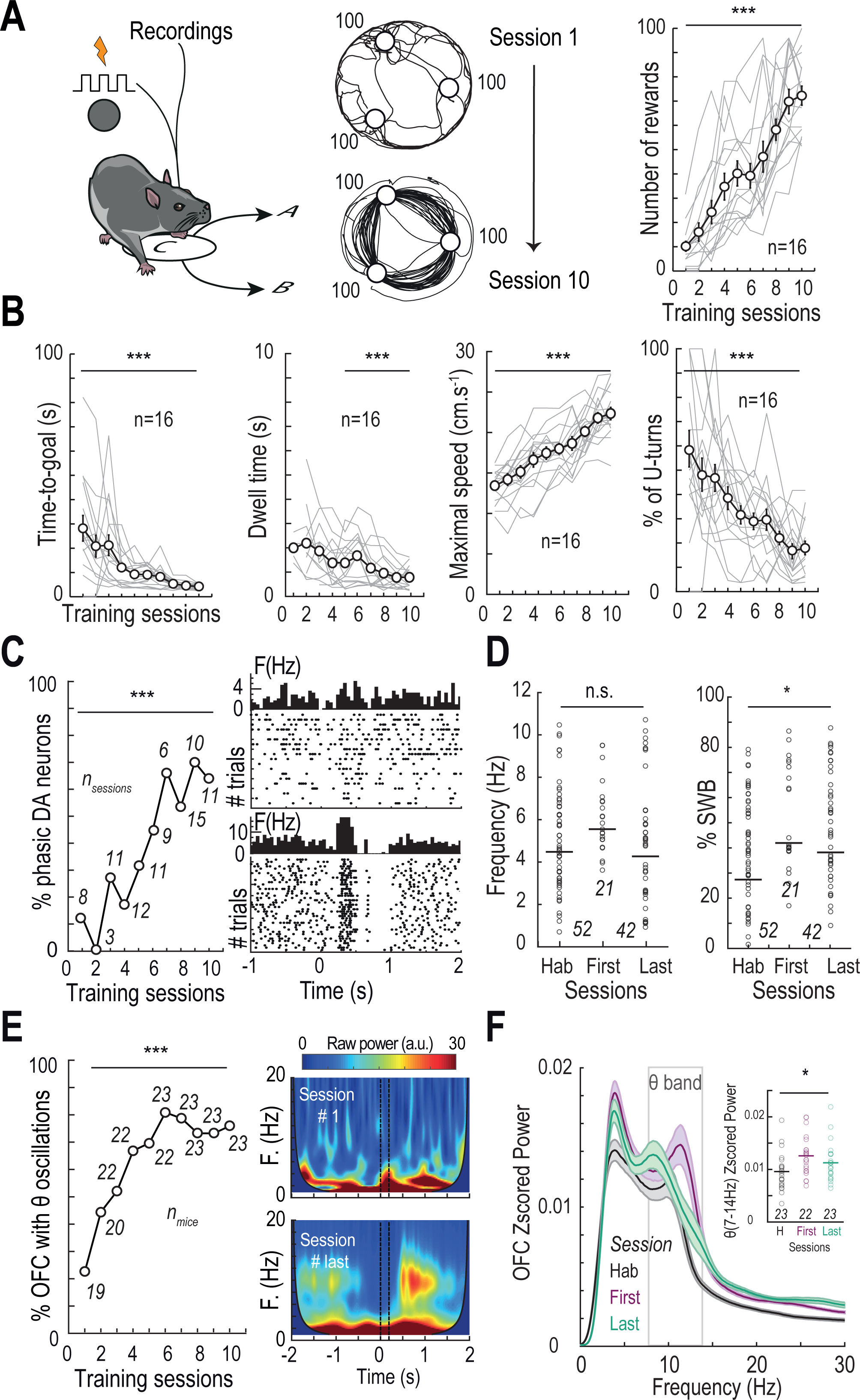
Early VTA and OFC activities emerge with learning. **(A)** Left: Schematic representation of learning in the D context with trajectory examples from session 1 (middle above) and session 10 (middle below). Right: Modifications of the number of rewards along learning. (Repeated measure ANOVA F_(9)_=45, p<0.001). Dots and vertical bar are mean ± SEM. Grey lines indicate modifications of the number of rewards per individual (n=13 animals) **(B)** Same as A for, from right to left: time to goal (repeated measure ANOVA, F_(9)_=11.9, p<0.001), dwell time (repeated measure ANOVA, F_(5)_=4.11, p<0.001), maximal speed (repeated measure ANOVA, F_(9)_=32.7, p<0.001), and proportion of U-turns (repeated measure ANOVA, F_(9)_=13.2, p<0.001) along the learning sessions. **(C)** Left: Proportion of pDAn modulated in-between two locations, along learning sessions (n=10) (χ^2^ test χ^2^=300, p<0.001). Right Examples of raster plots, centered on location entry, for a VTA pDAn early in learning (top) and another at the end of the learning (bottom) **(D)** Average firing frequency (left) and % spikes within bursts (right) throughout learning in the D context (Hab: open-field without ICSS prior to the learning stage, First: sessions 1-5, Last: sessions 6-10). Modification of firing frequency: ns ANOVA, F_(2)_=1.8, p=0.18; Same for %SWB: %SWB: ANOVA, F_(2)_=3.12, p=0.048. Dots and horizontal bar are mean ± SEM **(E)** Left: Proportion of OFC θ power (7-14 Hz) modulated in-between two locations, along learning sessions (χ^2^ test χ^2^=44.5 p<0.001). Right Examples of OFC time-resolved power spectral density centered on location entry, for one OFC early in learning (top) and for the same example OFC at the end of the learning (bottom). **(F)** Mean Zscored power of Fourier transform spectra of OFC during open field habituation (Hab, black, n=21, see Methods), early in learning (Det first, purple, n=20), at the end of the learning (Det end, green, n=19). Data are presented as mean ± SEM. Power in the θ-band frequency (7-14 Hz, grey box) shows a significant difference between the three conditions (ANOVA, F_(2)_=3.94, p=0.024), with differences between the Hab session and the first (Student t-test T_(43)_=2.80, p=0.015, Δ=+0.003) and last (paired Student t-test T_(22)_=-2.98, p=0.02, Δ=+0.002) sessions of the D context. No difference between first and lasts sessions (ns Student t-test T_(43)_=1.22, p=0.23).

This behavioral learning was associated with a modification of VTA pDAn dynamics. The proportion of pDAn with a firing rate significantly higher than baseline in-between two rewarded locations (see Methods) increased with the training sessions (Figure 3C left). Early in learning, when the behavior is still dominated by spontaneous locomotion, phasic pDAn firing was not locked to location entry (Figure 3C top right). In contrast, at the end of learning, increased pDAn activity appeared at similar timings after location entry across trials, as reflected in the significant increase when averaged (Figure 3C bottom right). However, the correlated locking to behavioral timings in dopamine firing during a trial, and the increase in the number of trials with learning, did not result in an overall (session wide) increase in pDAn firing frequency (Figure 3D left), but rather in a shift of pDAn firing pattern toward increased bursting (Figure 3D right). Hence early bursting in dopamine neurons did not rely on additional spikes, but rather on a dynamical re-organization towards bursting activity at behaviorally relevant times (Supp. Figure 3). Similarly, the total number of OFC EFPs in which θ oscillation power was higher than baseline in-between two rewarded locations gradually increased with learning (Figure 3E left). From relatively scarce task-locked oscillatory activity in θ band early during the learning, θ oscillations emerged consistently around the same timing after learning (Figure 3E right). When compared with power spectra from naïve animals undergoing a habituation session (Hab., Figure 3F) to the open field, θ power significantly changed throughout the learning of the task. More specifically, θ power increased in the first and in the last sessions of the deterministic context compared to naïve animals, but no difference was observed between the first and last sessions. Hence, θ oscillations occurred more often with learning and reorganized to lock at particular timings of the behavioral sequence. By contrast, no change in PFC oscillatory activity was observed throughout learning, which is expected as PFC does not display 8 oscillations at the end of the deterministic context (Figure 2). Overall, during the learning in the deterministic context, the optimization of trajectories and speed profiles were associated with a reorganization of VTA DA neurons firing towards time-locked bursting and of OFC activity towards time-locked θ oscillations.

### Bursting of VTA DA cells, together with frontal firing and oscillations, form a distributed signal for outcome discrepancy and expectations

Learning theories propose that behavior, and underlying brain activity, reorganize when unexpected outcomes occur: the discrepancy between observed and expected reward may be used to update the animals’ internal representations ^40–42^. This comparison is thought to be computed in mesocorticolimbic areas ^11, 43, 44^, most notably in VTA DA cells ^8^, but signals related to the evaluation of outcomes are also present in the frontal cortices ^45^. Yet, alternative interpretations, such as signaling outcome expectancy or prediction, may account for observed increases in neural activity associated with reward delivery ^46^. We thus evaluated VTA, OFC and PFC activities during expected rewards, and unexpected rewards and omissions. This further allowed to disentangle the direct effect of the intra-cranial stimulation on mesocortical activity (either as an artifact or as a reward response) from the reward representation learned by the animal. Early bursting activity in pDAn at the beginning of the probabilistic context occurred during the dwelling period and before the action initiation (Figure 4A top row, Supp. Figure 4A), as previously described for the end of the deterministic and probabilistic contexts (Figure 2). This increased DA activity was observed even when the ICSS reward was unexpectedly omitted at the beginning of the probabilistic context (Figure 4B top row, Supp. Figure 4A), but did not appear after unexpected, random stimulation reward in the home-cage (Figure 4C top row, Supp. Figure 4A). This suggests that, rather than being generated by the previous stimulation reward, early bursting at the beginning of the trial may be related to the expectation of the upcoming reward ^41, 47^, or the invigoration of the next movement ^21, 22^. The decreased pDAn activity following random stimulation in the home-cage emphasizes the emergence of early bursting activity in pDAn specifically upon self-paced actions. Moreover, pDAn activity decreased at the time of the expected reward during unexpected omissions (Figure 4B top row, Supp. Figure 4A), consistent with pDAn computing a reward prediction error ^7, 41, 48^.

**Figure 4:**
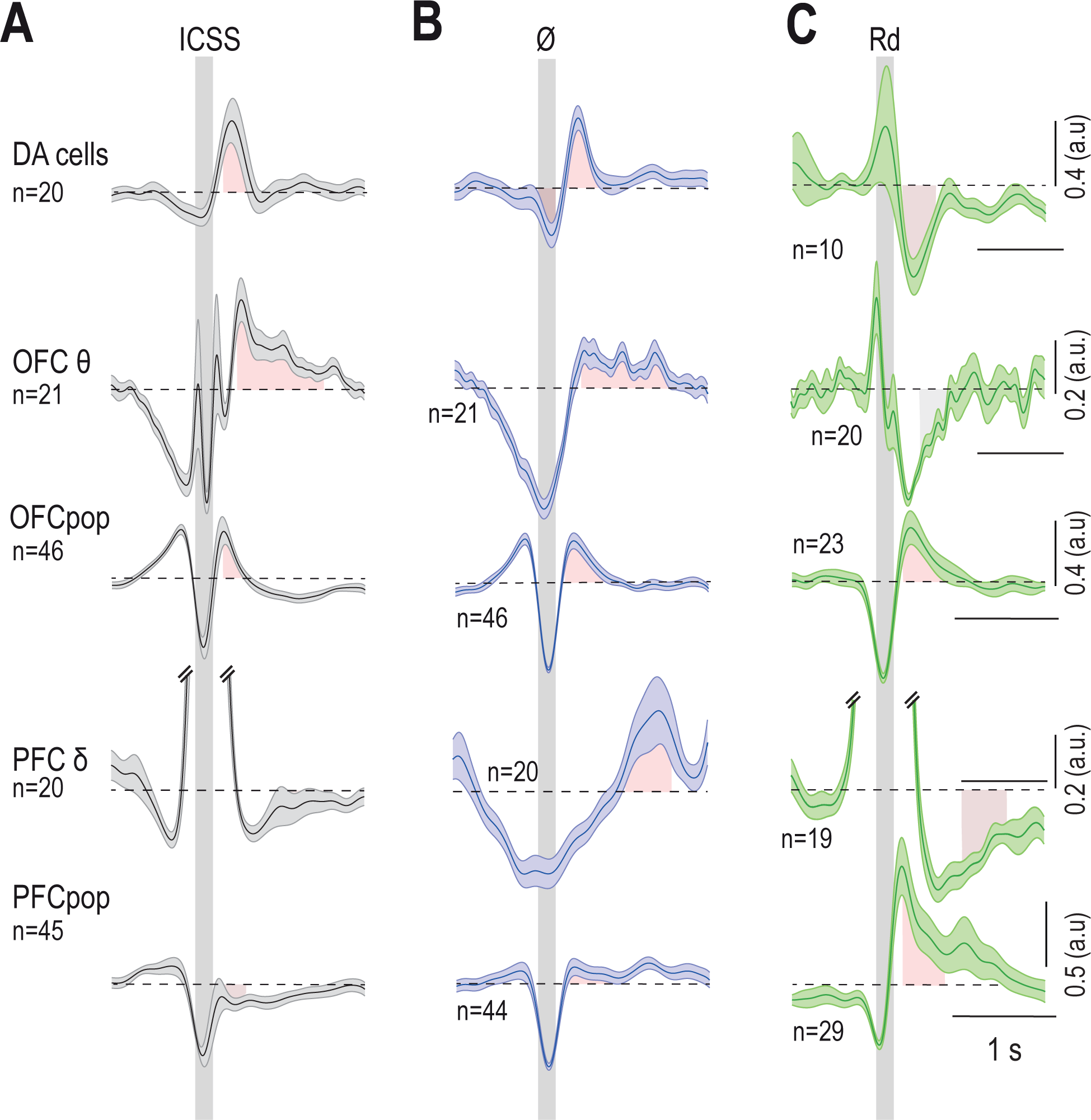
VTA, OFC and PFC activities following expected reward, unexpected omission and unexpected reward. **(A)** From top to bottom: VTA pDAn normalized firing, OFC θ oscillation power, OFC normalized population firing, PFC δ oscillation power, and PFC normalized population firing, centered on expected reward delivery upon location entry at the beginning of the P context. Data are presented as mean ± SEM. **(B)** Same as (A), centered on unexpected omission of reward delivery upon location entry at the beginning of the P context. **(C)** Same as (A), centered on unexpected reward delivery upon random intra-cranial stimulation in the home-cage, before the beginning of the conditioning. (Red) significant increases, (grey) ns activity, (brown) significant decreases.

Likewise, in the OFC (Figure 4 second and third row), neither the increased power of θ oscillations (Figure 4A second row, Supp. Figure 4B) nor the increased firing (Figure 4A third row, Supp. Figure 4B) at the beginning of the trials were triggered by the previous stimulation reward. Indeed, both θ oscillations and increased firing were observed during omission trials (Figure 4B, Supp. Figure 4B), and no difference was observed between the ICSS and omission conditions (Supp. Figure 4B left). Furthermore, unexpected, random stimulation rewards did not generate θ oscillations (Figure 4C, Supp. Figure 4B), indicating a specific involvement in self-paced behavior. Unexpected reward increased OFC population firing (Supp. Figure 4B right), suggesting an influence of both expected and unexpected outcomes on OFC firing activity.

Finally, in the PFC (Figure 4 fourth and last row), unexpected reward omission induced δ oscillations (Figure 4B forth row, Supp. Figure 4C left) and increased population firing (Figure 4B last row, Supp. Figure 4C right). Hence, δ oscillations and increased firing in the PFC, observed in the probabilistic context (Figure 2) but not in the deterministic context (Figure 4A), were already observed in the first omission trials. By contrast, unexpected stimulation reward decreased δ oscillations (Figure 4C fourth row, Supp. Figure 4C) and increased population firing (Figure 4C last row, Supp. Figure 4C), suggesting an involvement of the PFC specifically following unexpected outcomes (either reward or omission).

Overall, we observed in VTA, OFC and PFC firing and oscillations a distributed encoding of errors related to unexpected reward and omissions, which can be used for behavioral learning. Yet, these errors were distinct in each structure: VTA computed classical reward prediction error, PFC signaled unexpectedness while OFC firing signaled any event (unexpected or expected). Furthermore, we ruled out that the electrical stimulation was the direct cause of the observed mesocortical dynamics, as distributed activity was observed even after omission, but not after random ICSS.

### Distributed and complementary representations of decision parameters in mesocortical structures

The above analysis indicates that the mesocortical dynamics emerging with decisions were not only caused by the outcome immediately preceding. We thus searched for a signature of the internal representations of the upcoming outcome, which is thought to guide self-paced decisions ^1, 3^. First, we determined which of the task parameters actually affected the animals’ choices, by modeling how mouse choices depended on the reward probabilities of the locations and to the motor requirements of U-turns.

To do so, we extracted the succession of binary choices (as mice could not receive two consecutive rewards at the same location, they had to choose between the two remaining locations, Figure 5A left). We expressed choices in the probabilistic context as the proportion of exploitative choices in three gambles (G) between the potential outcomes (Figure 5A, G_25_:100 vs 50, G_50_: 100 vs 25 and G_100_ 50 vs 25% reward probabilities, respectively, exploitative choices consisting in choosing the option with the highest reward probability). Mice displayed a preference for higher reward probabilities in G_100_ (p=0.02) and G_50_ (p<0.001), but not in G_25_, in which they chose the locations associated with 50% and 100% reward in the same proportions (p=0.20). This replicated our previous studies ^32, 33^ in the probabilistic context: on average, mice assigned a positive value to uncertainty, which is zero for predictable outcomes (here, 100% probability) and maximal for the most unpredictable outcome (50% probability). As reward expectation (probability) and uncertainty (variance) co-vary in this setup, we used a model-based analysis to disentangle the influence of expected reward and uncertainty on choices. To quantitatively describe the decision processes underlying steady-state choice behavior in mice, we modeled individual data using a softmax model of decision-making ^32^. The model included: i) a value sensitivity parameter (β) measuring the trade-off between exploitative choices and random decisions, ii) a reward uncertainty bonus (ϕ) measuring how much animals value uncertain options and iii) a motor cost (κ) measuring the negative value of performing a U-turn ^31, 33^, as mice favored forward trajectories rather than U-turns (Figure 1), in both the deterministic and probabilistic contexts. We compared this model with simpler ones to assess the importance of the κ and ϕ parameters. Model comparison (Figure 5B) confirmed that in the probabilistic context, animals added the uncertainty to the expected reward as a total positive value on average, discounted by the motor cost (negative value) of U-turns (U model, Figure 5B).

**Figure 5:**
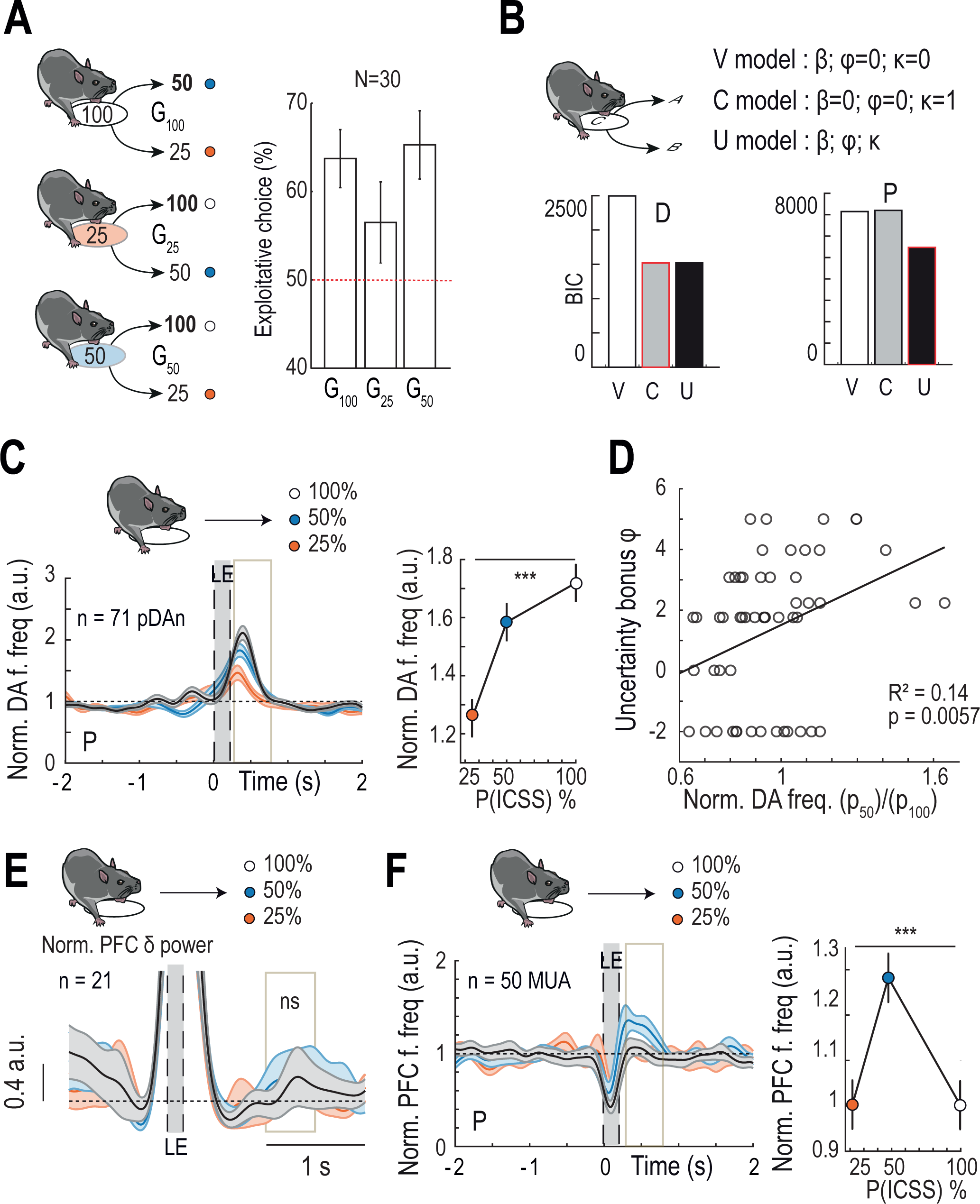
Complementary encoding of choices, value and cost by VTA, OFC and PFC. **(A)** Left: Scheme of the three gambles mice are facing in the probabilistic context. Right: Proportion of choices of the location associated with the highest reward probability for each gamble (G25: 100% vs 50%, G_50_: 100% vs 25% and G_100_: 50% vs 25%) at the end of the P context (N=30). Vertical bars represent SEM. **(B)** Bayesian Information Criteria (BIC) computed using three models of choice selection at the end of the D context (left) and of the P context (right). Red boxes surround the smaller BIC, indicating the best fit. V model: Softmax with β only (value sensitivity model), C model: Softmax with κ only (motor cost model), U model: Softmax with β, κ and φ (uncertainty bonus model) (See Methods). **(C)** Left: Normalized firing frequency (a.u.) of pDAn at the end of the P context, centered on location entries. Trials are sorted according to the chosen location. Grey box indicates the quantification window. Right: Quantification of the pDAn firing frequency according to the reward probability of the goal: one-way ANOVA, p<0.001. Data are presented as mean ± SEM. (**D**) Phasic encoding of uncertainty by pDA neurons (activity related to 50%, p_50_, versus 100%, p_100_, reward probability of the chosen locations) against the uncertainty bonus ϕ from the model (R^2^=0.14, p=0.006). **(E)** Same as (C) left for normalized δ (3-6 Hz) power (a.u.) of the PFC. (Mean δ power over 1-1.5s after LE according to the probability of the goal: ns ANOVA F_(2)_=0.07, p=0.93). **(F)** Same as (C) for normalized population firing frequency (a.u.) in the PFC. (Mean PFC firing over 0.3-0.8s after LE according to the probability of the goal: ANOVA F_(2)_=5.34, p=0.006).

We thus assessed the encoding of expected reward, uncertainty and motor cost in mesocortical activity to understand how outcome representations shapes self-direction. We did not find any encoding of motor cost in any of the recordings (Supp. Figure 5A). By contrast, VTA pDAn activity scaled with expectation of reward (Figure 5C), i.e. firing activity was minimal when the target location was p_25_ and maximal when going for p_100_. This finding is consistent with early bursting in pDAn signaling a quantitative “time-difference” reward prediction error ^41^. In this framework, DA cells not only signal the difference between actual and expected reward, but also integrate the expectation of future rewards predicted by the current state and actions of the animal ^8^. We further assessed whether DA cells only signaled the expected value of reward, or also integrated the bonus value of uncertainty. To do so, we computed the ratio between the encoding of the most uncertain option (p50, 50% probability) and the encoding of the most certain option (p_100_, 100% probability) by VTA pDAn activity, which indicates how much the encoding of the uncertain option deviates from a pure “reward expectation” signal. This p_50_/p_100_ ratio correlated with the uncertainty bonus parameter derived from animals’ decisions (Figure 5D), which measures how much an animal values uncertain options. This positive correlation between pDAn encoding of uncertainty and the value assigned to uncertain options suggests that VTA pDAn integrate uncertainty with reward into a common currency ^49^, thereby promoting the choice of the uncertain options ^32^.

PFC θ oscillations power did not scale with the expectation of reward uncertainty (Figure 5E), in contrast to PFC population firing (Figure 5F) that was maximal when mice moved towards the location associated with the most uncertain (50%) reward probability. We further found that PFC population activity was enriched with the encoding of reward uncertainty above chance level (Supp. Figure 5B). Hence, while PFC δ oscillations reflected preceding unexpected outcomes (Figure 4), population firing signaled the expected uncertainty of the predicted outcome. Finally, we did not find evidence for OFC θ oscillations (nor firing) to encode expected reward or uncertainty (ns ANOVA F_(2)_=0.3, p=0.97). OFC oscillations were uncorrelated to the expectation of outcome value. They occurred following both reward and omission during the task, but not after passive exposition to unexpected rewards in the home-cage (Figure 4). Together, this suggests an OFC specificity related to active behaviors, rather than to the expectation of outcomes. Therefore, the internal representations of reward outcomes influencing animals’ self-directions, i.e. expected reward and uncertainty, were represented in a complementary way by the VTA and the PFC activities, respectively, while the OFC encoded a more general, active change in the behavioral state of the animals.

### Distributed correlates of self-initiation, invigoration and pace of goal-directed actions

After examining the internal representations directing self-paced behavior, we then assessed the relation between circuit activity and the content of self-paced actions. We analyzed how the initiation and invigoration of goal-directed behaviors related to mesocortical dynamics at each trial. As shown above (see Figure 1), each trial is characterized by the initiation of a movement towards the next location (estimated by the dwell time) and a strong acceleration (estimated by the time to maximal speed) followed by a deceleration at the next location, resulting in the overall time to goal (see Figure 1, Figure 6A top). Then another trial ensues. Sorting the trials by ascending pDAn early bursting for each neuron revealed a negative correlation with both the time to goal (Figure 6A) and the time to reach the maximal speed (Figure 6B-C): greater pDAn phasic activity before the self-initiation correlated with shorter time to goal, due to a shorter time to reach the maximal speed. This was confirmed by the distribution of correlation coefficients (R^2^) for all neurons (Figure 6D). We ruled out that these correlations might have been spurious using surrogate data preserving the coding capacity of pDAn cells and the timing variability of mice (Figure 6D, see Methods).

**Figure 6:**
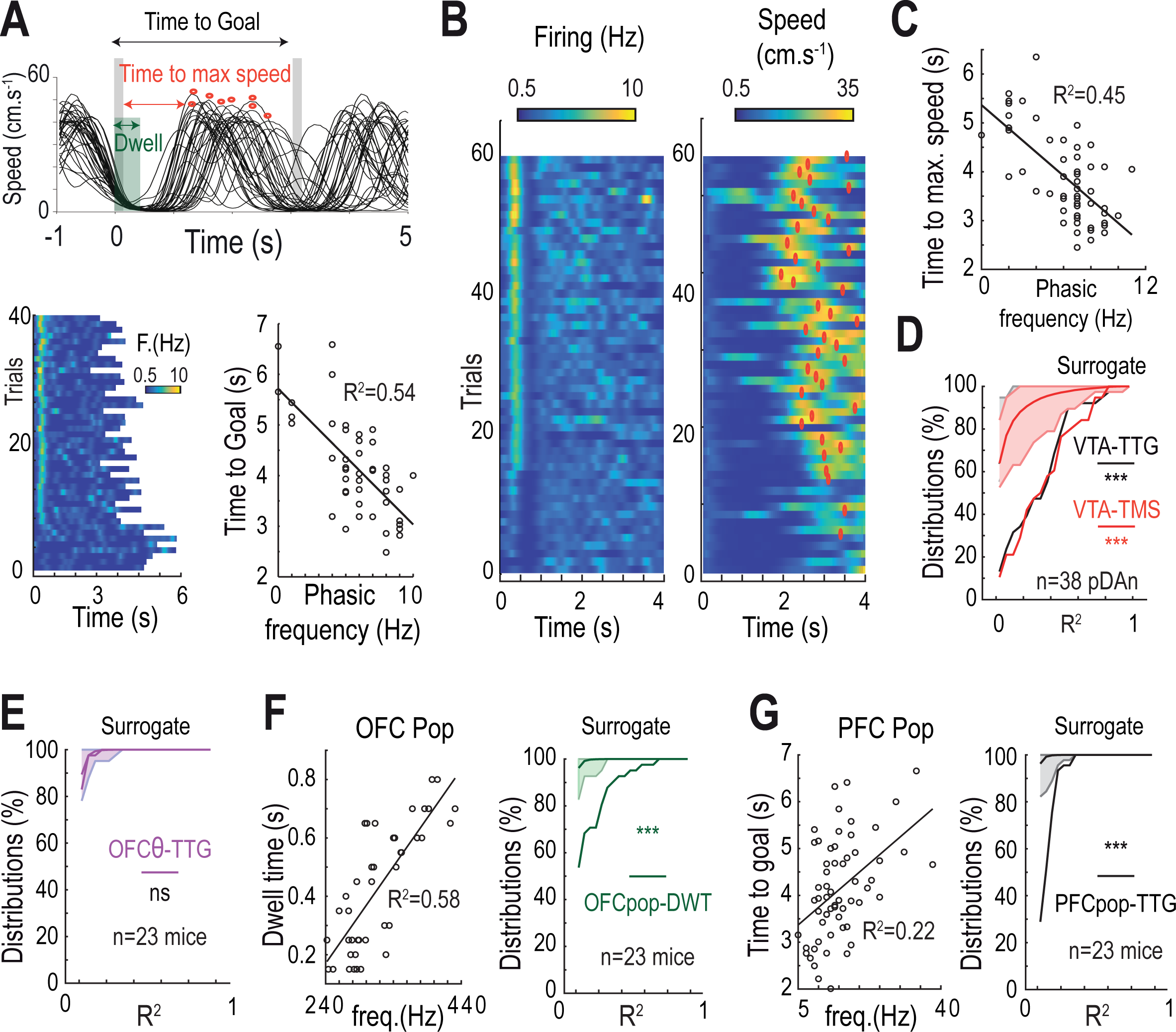
Distributed encoding of self-initiation, invigoration and pacing by OFC VTA and PFC. **(A)** Top: Example of instantaneous speed profile for one mouse in the D context. Grey boxes indicate ICSS durations, the green box the dwell time and red dots the maximal speed within trials. Bottom: Example of firing frequency for a pDAn in the D context with trials sorted from the smallest to the highest bursting frequency after trial initiation (left), and relation between the time to goal and the frequency of the early bursting activity (right) each dot representing a trial. **(B)** Example of firing frequency for another pDAn in the D context with trials sorted from the smallest to the highest bursting frequency after trial initiation (left), and the instantaneous speed profile associated for each trial (right), red dots indicating the maximal speed within trials. **(C)** Relation between the time to maximal speed within trials and the phasic frequency for the cell showed in (B). **(D)** Distribution of correlation coefficients (R^2^) for correlations between phasic frequency (Hz) of pDAn and time to goal (black) and time to maximal speed (red) at the end of the D context for n=38 cells. Data are presented as mean ± SEM. Surrogate data are generated by computing correlations with shuffled firing frequency and time to goal (or time to maximal speed). Kolmogorov-Smirnov test of actual distribution versus surrogates: p<10^-3^ for time to goal and time to maximal speed. **(E)** Distribution of correlation coefficients (R^2^) for correlations between OFC θ (7-14 Hz) power and time to maximal speed at the end of the D context. Data are presented as mean ± SEM. Surrogate data are generated by computing correlation with shuffled θ power and time to maximal speed. Difference with surrogates: ns Kolmogorov-Smirnov test p=0.62. **(F)** Left: Example of relation between the dwell time and the population firing frequency of OFC of one mouse at the end of the D context, each dot representing a trial. Right: Distribution of correlation coefficients (R^2^) for correlations between OFC population firing frequency and dwell time. Data are presented as mean ± SEM. Surrogate data are generated by computing correlation with shuffled firing frequency and dwell time. Difference with surrogates: Kolmogorov-Smirnov test p<10^-3^. **(G)** Same as (F) for PFC population firing frequency and time to goal. Difference with surrogates: Kolmogorov-Smirnov test p<10^-3^.

By contrast, we did not find any significant correlation between the amplitude of cortical oscillations and the successive behavioral timings, neither for OFC θ nor for PFC δ oscillations (Figure 6E, Supp. Figure 5C-F). This might be due to the temporal order of neural oscillations and behavioral events: the increase in OFC θ and in PFC 8 generally occurred after self-initiation (see Figures 2 and 4), and thus may not be involved in this decision process. On the opposite, the increase in OFC population activity, which occurred early in the trial, correlated positively with the dwell time, suggesting an involvement in the self-initiation of the trial (Figure 6F). Finally, PFC population firing correlated with the time to goal, but neither with the dwell time nor with the time to maximal speed (Figure 6G). This indicate that additional variability in the overall pace of the trial, that was not already due to earlier decisions (Figure 1), may be encoded in the PFC. Overall, OFC, VTA and PFC firing activity synergistically encoded for self-initiation, invigoration, and pace of the goal-directed actions, suggesting a sequential and distributed mechanism for self-paced decisions.

### Synergy and antagonism between mesocortical structures in self-paced decisions under uncertainty

Electrophysiological analyses suggest an involvement of the VTA and PFC in both the selection (Figure 5) and execution of actions (Figure 6). We thus investigated their causal involvement, by inactivating these structures during the task. To specifically manipulate VTA DA neurons, we expressed an inhibitory halorhodopsin variant (Jaws ^50^) in DAT^iCRE^ mice using a Cre-dependent viral strategy (Figure 7A). We confirmed expression of the opsin in Jaws-transduced mice with immunochemistry and verified that 500ms-light pulses (520 nm) at 0.5 Hz reliably decreased the activity of VTA DA neurons using patch-clamp recordings (Sup Figure 6A). As the amplitude of the early bursting in VTA pDAn correlated with movement invigoration (Figure 6), we specifically tested the effect of optogenetic inactivation of VTA DA neurons at the time of the early bursting activity (using 500ms continuous light, starting 100ms after the previous ICSS, see Methods, Figure 7A). Optogenetic inhibition on each trial in the deterministic context decreased the total number of transitions (Figure 7B). The decrease in the number of trials was due to an alteration in the speed profile that started during illumination and lasted after its termination (Figure 7C), with an increase in the overall time to goal (Figure 7D left). In particular, photo-inhibition of VTA pDAn delayed the action initiation (increased dwell time, Figure 7D middle) and decreased the action vigor (decreased maximum of mean speed, Figure 7D right). None of these parameters were modified in GFP-transduced DAT^iCRE^ mice (Supp. Figure 6B). These results causally implicate VTA DA cells in motivation for ongoing, self-generated movements, energizing movement as previously observed ^21, 47^, but also promoting movement initiation ^22^.

**Figure 7:**
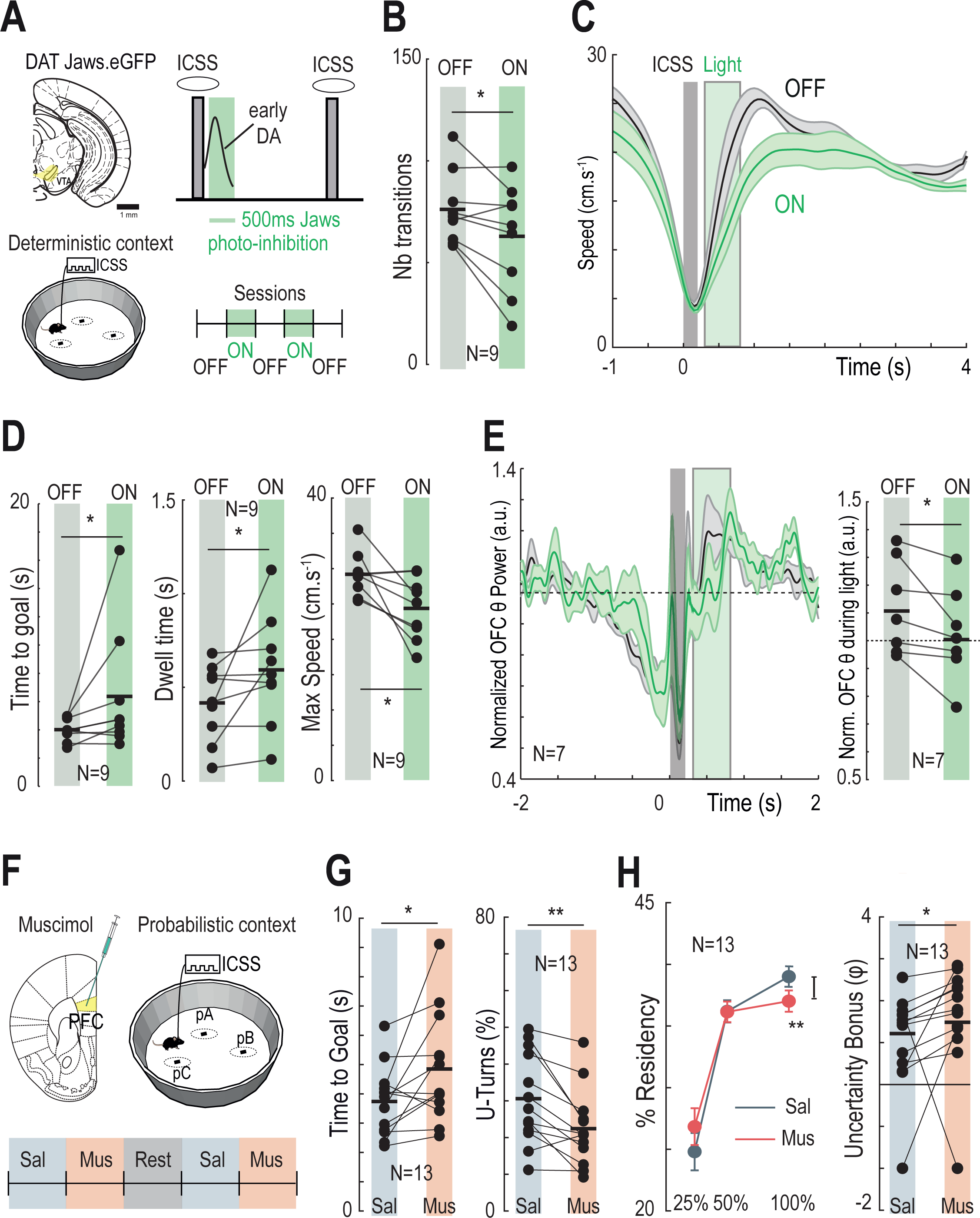
Behavior-dependent synergy or antagonism between DA VTA neurons and frontal cortices. **(A)** Schematic representation of the manipulation experiment using the inhibitory opsin Jaws. Jaws is expressed unilaterally in the VTA of DATi^CRE^ mice using CreLox strategy. The light was applied continuously for 500 ms, 100 ms after the ICSS (see Methods), in order to suppress the phasic pDAn activity. Mice underwent at the end of the D context, a succession of OFF (light OFF) and ON (light ON) sessions. **(B)** Effect of ON light stimulations compared to OFF on the number of transitions (*i.e.* of rewards) obtained (paired Student t-test T_(8)_=2.54, p=0.03, Δ=-13.2). Horizontal bars represent means. **(C)** Effect of OFF or ON light stimulations on instantaneous speed profile within trials. Data are presented as mean ± SEM. Grey box indicates the ICSS duration and green box the light duration. **(D)** Effect of ON light stimulations compared to OFF on the time to goal (left), the dwell time (middle) and the maximal speed (right). (Effect on the time to goal: paired two-sided Wilcoxon signed rank test W_(8)_=5, p=0.04, Δ=2.40s; Dwell time: paired two-sided Wilcoxon signed rank test W_(8)_=3, p=0.0195, Δ=0.17s; Max. speed: paired Student t-test T_(8)_=3.04, p=0.016, Δ=-4.81 cm.s^-1^). Horizontal bars represent means. **(E)** Left: Effect of ON light stimulations compared to OFF on the normalized OFC θ power (7-14 Hz) (paired Student t-test T_(6)_=2.63, p=0.039, Δ=-0.097 a.u.). Data are presented as mean ± SEM. Grey box indicates the ICSS duration and green box the light duration. Right: Quantification of OFC θ power during light duration. Horizontal bars represent means. **(F)** Schematic representation of the PFC inactivation experiment using bilateral muscimol infusion at the end of the P context. Mice underwent a succession of sessions following saline or muscimol infusions. **(G)** Effect of muscimol on the time to goal (left) and the proportion of U-turns (right) compared to saline. (Time to goal: paired two-sided Wilcoxon signed rank test W_(12)_=12, p=0.0171, Δ=+1.11s; U-turns: paired Student t-test T_(12)_=3.79, p=0.0026, Δ=-8.18%). Horizontal bars represent means. **(H)** Left: Effect of muscimol on the choice repartition between the three locations compared to saline. (Effect on the p_100_ choice: paired Student t-test T_(12)_=3.71, p=0.003, Δ=-1.99%). Data are presented as mean ± SEM. Right: Effect of muscimol, compared to saline, on the fitted φ parameter (uncertainty bonus) obtained using the Softmax based on three parameters β, φ and κ (see Methods) (paired two-sided Wilcoxon signed rank test W_(12)_=13, p=0.0215, Δ=+0.54). Horizontal bars represent means.

VTA DA cell photo-inhibition also affected OFC θ oscillations (Figure 7E left), with a decrease in θ oscillations power during photostimulation (Figure 7E right) which did not last afterwards. We did not find any effect of VTA DA cell photo-inhibition on other frequencies of the OFC power spectra, nor in PFC oscillation, suggesting a specific relation between pDAn early bursting and OFC θ oscillations. As the OFC dynamics transitioned from increased population firing to θ oscillations at action initiation (Figure 2C, Figure 7E), VTA DA neuron photo-inhibition may have delayed θ oscillations directly, or indirectly, with OFC θ oscillations merely following the delayed action initiation. This latter interpretation is consistent with the peak in OFC θ power occurring at 0.62s (and a dwell time of 0.41s) without light; and 0.83s (with a dwell time of 0.59s) under photo-inhibition.

The absence of observable effects of VTA DA cell photo-inhibition on PFC dynamics is also in line with our electrophysiological results suggesting that the PFC is not implicated in self-initiation in the deterministic context. In the P context however, PFC δ oscillations have been specifically detected, and PFC firing frequency has been correlated with time to goal and reward uncertainty. We thus probed the involvement of the PFC in the P context by inactivating this structure bilaterally using a local infusion of muscimol (Figure 7F, Supp. Figure 6C).

PFC inactivation in the probabilistic context increased the time to goal (Figure 7G left), which was associated with an increase of the dwell time and a decrease in maximal speed (Supp. Figure 6D). This resulted in a lower number of transitions (Supp. Figure 6D). These results confirm our electrophysiological data showing an encoding of time-to-goal by PFC population firing (Figure 5), and suggest a role for the PFC in the overall pace of the action.

Surprisingly, muscimol also decreased the percentage of U-turns (Figure 7G right). This was due to an alteration of choices: PFC inactivation affected choice repartition on the three locations (Figure 7H left) with a decreased propensity to visit the location associated with the highest reward probability (i.e. 100%). We thus sought to explain these changes in U-turns and in the choices of the best locations using the computational model. We fitted the transition function of each mouse (Supp. Figure 6E) in the saline and muscimol sessions with a model taking into account the value sensitivity (β), the weight of uncertainty (φ) and the motor cost (κ) of each decision (see Methods and Model U, Figure 5B). We found that under muscimol, animals behaved as if their uncertainty bonus was amplified (Figure 7H right), with no changes in nor in κ (Supp. Figure 6F). Again, PFC inactivation with local muscimol extends our electrophysiological results: PFC population firing positively encoded reward uncertainty, and PFC inactivation increased the valuation of uncertainty, suggesting an inhibitory control of the PFC on uncertainty-seeking in our setting.

## Discussion

By using a task in which mice perform stereotyped trials from a motor perspective (moving from one rewarding location to another), we eliminated the requirement for a stimulus to reiterate and time-lock a specific behavior, usually needed for statistical analyses. Furthermore, the absence of a cue specifying the direction or the initiation of each trial produced variability (mice had to choose among several alternatives and to decide the pace of goal-directed actions) enabling correlation of behavior with neural activity. This allowed us to assess how self-generated actions arise from the contextual reorganization of mesocortical dynamics. We found that a distributed sequence of firing and oscillations in the VTA, PFC and OFC jointly set the goal of actions (i.e., deciding where to go), self-initiated the trial and determined the vigor and pace of the goal-directed action. This circuit sequence was influenced by the reward context (deterministic or uncertain) and correlated with reward value and uncertainty, used to guide the animal’s choices. Both the cortical oscillations and the distributed increase in firing emerged during learning as a reorganization of existing dynamics, and all these structures encoded prediction errors about the outcomes. Such sequence, rather than being fixed, could incorporate the PFC or not, depending on the reward context, and the PFC could act in synergy or antagonistically to the VTA in co-determining action selection or execution.

### Modular or distributed neural encoding of decisions

A long-standing question in decision-making remains to identify the elements of the mesocorticolimbic circuits with computation underlying decisions. Many theories assume that goal-directed choice can be subdivided into a set of localized modules, each computing a variable of a phenomenological model of behavior ^6, 15^. For instance, in the influential reinforcement-learning framework, the phasic activity of dopamine neurons is classically viewed as calculating a reward prediction error (RPE, ^8^), i.e., the difference between actual and predicted value. In this framework, prefrontal and orbitofrontal cortices are implicated in value predictions, task state inference or beliefs ^10, 24, 39^. In stimulus-driven behaviors, frontal cortices are thus considered as upstream computations to dopamine RPE, inferring value or state predictions to be compared with actual outcomes by DA cells, while the frontal areas play an indirect role in behavior, by modulating a learning signal ^3, 9, 10, 24, 39^ or by informing areas that perform the choice, typically the basal ganglia ^3, 9, 39^. By contrast, pharmacological and chemogenetic studies have shown an implication of prefrontal dopamine in decision-making, in particular under uncertainty ^27, 51, 52^, placing cortical dynamics downstream of dopamine modulation. This is in line with a vast literature assigning a more direct role to frontal cortices in ongoing action selection ^14, 15, 53^. The OFC and PFC have been shown to play crucial roles in ongoing reward-directed choice. In particular, neurons in the OFC have been proposed to encode the subjective value of options and to bias future choice ^11, 54–56^ while neurons in the PFC have been suggested to compare the values of options in order to implement action policy ^6, 14, 53^. In particular, θ oscillations, widely observed across frontal cortical areas, are also implicated in general functions such as reward processing and goal-directed behavior^16,52,57,58^.

Our recordings of neural activity in the VTA, the OFC and the PFC during the task revealed a distributed sequence of neural activity, which was consistent in each trial. This distributed mesocortical sequence started with an early phasic activity in the DA system together with an increase in OFC firing, before the initiation of the action. This self-initiation was followed by a transition from increased spiking in the OFC to a θ oscillation. Increased OFC firing encoded the time of self-initiation, which may suggest an early cortical participation in decision-making. However, using optogenetic inhibition, we further show that VTA DA cells are likely to emit the “go” signal for initiation (see also below) as well as to favor the transition from desynchronized spiking to θ oscillation in the OFC. Our results thus suggest a recurrent OFC-VTA-OFC computation related to action initiation. We also found lines of evidence for a distributed computation of the overall pace of the action (from one rewarded location to another) among the PFC and VTA: firing in both structures correlated with the time to goal, which was also increased by the local inhibition of each structure. Our study is therefore in line with the alternative perspective that decisions may be distributed, i.e. formed in concert across the subparts of the mesocorticolimbic loop that perform parallel computations ^6, 15, 59^. The same view may hold for learning, as errors about the outcomes were present in all structures, during unexpected reward or omissions. Nevertheless, prediction errors were not similar: while VTA firing did conform to a reward prediction error (decreasing with omission and increasing with reward), the OFC signaled both (expected and unexpected) outcome regardless of its valence, and the PFC was engaged by unexpectedness only, in particular during omissions. Hence, for the learning part, our results are consistent with reinforcement-learning views that the VTA relates to value prediction errors ^8, 60^ while the OFC is more dedicated to detect state changes ^11, 39, 43, 61^ and the PFC to uncertainty ^10^ or deliberative strategies when the environment is uncertain ^62–65^. Such complementary predictions errors in the VTA and frontal areas may arise from the same kind of comparisons ^6^ applied to different inputs, and might be needed for learning in complex settings ^39, 43, 66^. More studies are thus needed to reconcile the modular and distributed views of learning and decision-making, with a potential important role for biophysical models to integrate neurophysiological findings at the circuit level ^6^.

### Roles of mesocortical circuits in ongoing decisions

In recent studies, it remains debated whether DA RPE at action initiation constitutes a learning signal for the next trials ^20, 67^ and/or a motivational command for the current trial ^19, 22, 47, 68^. In our recordings, we observed that early bursting in VTA DA neurons was not directly triggered by the MFB electrical stimulation. Indeed, early VTA DA bursting scaled with the reward probability of the future location, and it decreased at the time of the expected reward during reward omissions. All of these results comply with early bursting in VTA DA cells encoding RPE. However, early VTA DA bursting also correlated with the vigor of the movement (estimated by the time to maximal speed and the time to goal). Furthermore, inhibition of this phasic activity reduced the time to goal by decreasing the maximal speed and delaying the action initiation. We thus causally implicate RPE-complying VTA DA activity in the initiation and vigor of goal-directed actions ^19, 22^, contrary to the view that VTA DA activity only constitutes a passive reflection of action initiation ^20^. As pDAn firing did not encode the time of initiation in advance, and because temporarily inhibiting VTA DA cells delayed initiation, we propose that VTA DA activity participates to a “go” command ^22, 68^, with computations leading to action initiation performed elsewhere, in the OFC based on the present data, but also in the striatum ^3, 15^. The encoding of dwell time by the OFC, as well as the transition from increased spiking to θ oscillations at action initiation, are in line with the state theory of the OFC ^39, 61^. In this theory, the OFC computes a “you are here” signal (within the task space), which is based on both external (cues) and inferred information. Action initiation would constitute a change in the animal’s state, associated with distinct OFC dynamics. This may also explain why OFC firing and oscillations react to whichever outcome (reward or omission), to signal changes between the different possible states of this task. However, in our task, action initiation is not caused by external cues signaling a state change, nor by the inference of a hidden state, such as for instance an non-signaled yet imposed delay after which the action has to be initiated ^10^. On the opposite, we found that the OFC computed in advance the duration before state change (i.e. the dwell time), suggesting an active involvement of this structure in controlling the change between task states (a “you wait here” signal), rather than a passive role in monitoring task states.

In line with a general role related to task space, we did not see any correlation between the discharge of OFC neurons, or the amplitude of OFC θ oscillations, and the expected reward nor the reward uncertainty. This is at odds with accounts suggesting involvement of the OFC in economic value ^14^ or in computing confidence (i.e., the inverse of uncertainty, ^69, 70^). However, value or uncertainty can be confounded with arousal and salience and causal involvement of the OFC in economic choice is lacking ^66^. In contrast with the OFC, we found clear correlates of value and uncertainty in the VTA DA cell firing, and of uncertainty in the PFC firing. Quantitative encoding of probabilistic reward complies with RPE theories and has been described at length ^71–73^. PFC δ oscillations have also been implicated in motivation ^74^. However, expected uncertainty was not aggregated with expected value in PFC activity, contrary to observations made in humans ^75, 76^. This might relate to task differences rather than species differences. Indeed, the PFC has been implicated in selecting the strategy for goal-directed choice ^16, 39^. In our task, we did not observe any increased firing nor any δ oscillations in the PFC in the deterministic context, suggesting that the PFC is mostly needed in the probabilistic context, i.e. for decisions under uncertainty ^62, 64^. As mice used uncertainty to guide their decisions in the probabilistic context, the PFC may encode uncertainty in our task in order to represent decision-guiding rules, rather than uncertainty-modulated values per se.

### Context-dependent synergy or antagonism between PFC and VTA

Inhibiting the PFC by a local infusion of muscimol led to an increase of uncertainty-seeking, indicating that the encoding of uncertainty by the PFC exerts an inhibitory influence on uncertainty-biased choices. As dopamine generally exerts a positive influence on uncertainty-seeking ^32, 51^, this suggests antagonistic influences of the VTA and PFC on the motivation induced by reward uncertainty. This might reflect a difference between how model (or belief)-based and model-free control may treat uncertainty. In the probabilistic context, the uncertainty associated with the reward is known by the animals, i.e. it is a form of expected uncertainty ^77^. In the simplest form of curiosity, expected uncertainty or variability may exert a positive motivational influence in the form of a bonus added to the expected value by DA cells, in order to promote the exploration of unpredictable options ^32, 78^. By contrast, in deliberative strategies putatively implicating the PFC, the known, expected uncertainty may be treated as uninformative noise that has to be discarded from the decision strategy ^77^. Hence, expected uncertainty may be incorporated by VTA DA neurons into value to promote model-free exploration, and encoded by PFC to favor model-based exploration, thus opposing model-free uncertainty-seeking.

Strikingly, the VTA and PFC had opposite influences on uncertainty-related choices, but synergistic influences on the pace of actions. Suppressing the early phasic activity of the pDA neurons and inhibiting the PFC had similar consequences on the pace of behavior, with an increase in time to goal and a decrease in total transition number. This could either suggest a sequential computation (e.g. DA-then-cortex), a recurrent one (DA-cortex-DA, as suggested above for the VTA and OFC), or two independent processes causally linked to movement initiation. More work is thus needed to dissect PFC-DA interactions ^23, 26, 30^ in ongoing, self-generated decisions. We however suggest that this circuit presents a “fluid” organization as described for invertebrate circuits ^79^: depending on the reward context, the PFC may flexibly integrate the basic circuit formed by the VTA-OFC loop in charge for the initiation of a goal-directed action, adding uncertainty-based computations to the distributed sequence. In the same vein, the emergence of these activities with learning followed a reorganization of existing circuit dynamics. Indeed, completion of learning induced the emission of bursts by dopamine cells at each trial, and was also associated with an increase in the number of trials. The combination of these two features could have resulted in an overall increase in pDAn firing frequency, but we did not observe any change. Hence early bursting in dopamine neurons did not rely on additional spikes, but rather on a dynamical re-organization towards time-locked bursting activity. This increased in early DA activity correlated also with a locking of θ oscillations in the OFC. This suggests that the characteristics of the VTA-OFC-PFC, i.e. distributed yet distinct contributions to both learning and decisions, may rely on the alignment of distributed dynamics at relevant timings, rather than on increases in activities of separate modules each computing a decision variable. While this alignment of mesocortical dynamics is usually forced by a stimulus, we show here that it reorganizes even without stimulus, resulting in the self-generation of goal-directed actions.

## Acknowledgements

We are grateful to the animal facilities (IBPS), Camille Robert and Paris Vision Institute AAV production facility for viral production and purification. This work was supported by the Centre National de la Recherche Scientifique CNRS UMR 8246, INSERM U1130, the Foundation for Medical Research (FRM, Equipe FRM DEQ2013326488 to P.F), the French National Cancer Institute Grant TABAC-16-022 et TABAC-19-020 (to P.F.), French state funds managed by the ANR (ANR-16 Nicostress to PF).

## Author contributions

JN and PF designed the study. EB SD ST CPS MC JN performed the electrophysiological recordings. EB YAT CPS performed some behavioral experiments. SD and ST contributed to setup developments. EB JN SD ST CPS performed the surgeries and virus injections. EB performed the immunohistochemistry. LT provided the DAT-Cre mice. AM developed the optogenetics setup. JN and PF developed the model. JN, EB and PF analyzed the data. JN, EB and PF wrote the manuscript with inputs from the other authors.

## Methods

### Animals

Experiments were performed on adult C57Bl/6Rj DAT^iCRE^ and Wild-Type (Janvier Labs, France) mice. Male mice, from 8 to 16 weeks old, weighing 25-35 grams, were used for all the experiments. They were kept in an animal facility where temperature (20 ± 2°C) and humidity were automatically monitored and a circadian light cycle of 12/12-h light-dark cycle was maintained. All experiments were performed in accordance with the recommendations for animal experiments issued by the European Commission directives 219/1990, 220/1990 and 2010/63, and approved by Sorbonne University.

### AAV production

AAV vectors were produced as previously described using the cotransfection method and purified by iodixanol gradient ultracentrifugation^51^. AAV vector stocks were tittered by quantitative PCR (qPCR)^52^ using SYBR Green (Thermo Fischer Scientific).

### Intracranial self-stimulation electrode and recording electrode implantation

Mice were anaesthetized with a gas mixture of oxygen (1 L/min) and 1-3 % of isoflurane (Piramal Healthcare, UK), then placed into a stereotaxic frame (Kopf Instruments, CA, USA). After the administration of a local anesthetic (Lurocain, 0.1 mL at 0.67 mg/kg), a median incision revealed the skull which was drilled at the level of the Median Forebrain Bundle (MFB), the OFC, the PFC or the VTA. Dental cement (SuperBond, Sun Medical) was used to fix the implant to the skull. A bipolar stimulating electrode for ICSS was then implanted unilaterally (randomized) in the brain (stereotaxic coordinates from bregma according to mouse after Paxinos atlas: AP -1.4 mm, ML ±1.2 mm, DV -4.8 mm from the brain). Bipolar recording electrodes were implanted in the lateral OFC (AP +2.6 mm, ML ±1.5 mm, DV -1.7 mm from the brain) and the medial PFC (AP +1.65 mm, ML ±0.5 mm, DV -1.8 mm from the brain). Multi-electrodes were implanted in the VTA (AP -3.15 to -3.25 mm, ML ±0.5 mm, DV -4.1 to 4.25 mm from the brain). After stitching and administration of a dermal antiseptic, mice were then placed back in their home-cage and had, at least, 5 days to recover from surgery. An analgesic, buprenorphine solution at 0,015 mg/L (0.1 mL/10 g), was delivered after the surgery and if necessary, the following recovering days. The efficacy of electrical stimulation was verified through the rate of acquisition during the deterministic context (see behavioral methods).

### Virus injections

DAT^iCRE^ mice were anaesthetized (Isoflurane 1-3%) and were injected unilaterally (randomized left/right side and ipsi/contralateral side) in the VTA (1 µL, coordinates from bregma: AP -3.15 to -3.25 mm; ML ±0.5 mm; DV -4.55 mm from the skull) with an adeno-associated virus (AAV5.EF1α.DIO.Jaws.eGFP 1.16e^13^ ng/µL or AAV5.EF1α.DIO.YFP 6.89e^13^ or 9.10e^13^ ng/µL). A double-floxed inverse open reading frame (DIO) allowed to restrain the expression of Jaws (red-shifted cruxhalorhodopsin) to VTA dopaminergic neurons.

### Polyelectrodes

Hand-made multi-electrodes (2 bundles of 8 electrodes) were obtained by twisting eight polyimide-insulated 17 µm Nickel-Chrome wires. The use of eight channels relatively close together allows for a better discrimination of the different neurons. Before implantation and recording, the multi-electrodes were cut at suitable length and plated using a Platinium-PEG solution to lower their impedance to 150-400 KOhms and improve the signal-to-noise ratio. The free ends of the multi-electrodes were connected to the holes of EIB-18 (electrode interface board, Neuralynx) and fixed with pins. We manufactured a microdrive system (home-made 3D conception and printing) consisting of a main body, on which is mounted the EIB, and a driving screw, with a sliding part design to contain the two multi-electrodes. This microdrive allowed moving through the VTA in order to sample neuronal populations.

### Bipolar electrodes

Hand-made bipolar electrodes were obtained by twisting two Teflon-insulated (60 µm) Stainless Steel wires. Two configurations were used. For the first one, the tips of the bipolar electrodes were cut so that they are spaced of less than 0.5 mm apart. For the second one, the reference tip was wound around the recording one, at a distance of less than 0.5 mm from the recording endpoint. These electrodes are designed so the two tips are oriented perpendicular to the dipoles formed by cortical pyramidal neurons. The first configuration was used for OFC recording electrodes, and the second one for PFC recording electrodes. IntraCranial Self-Stimulation (ICSS) electrodes were made as the second configuration with an 80 µm Stainless Steel wire. Bipolar electrodes were connected to the EIB during the surgery, by fixing the free ends with pins.

### Immunochemistry

After euthanasia, brains were rapidly removed and fixed in 4% paraformaldehyde (PFA). After a period of at least three days of fixation at 4°C, serial 60-µm sections were cut with a vibratome (Leica). Immunostaining experiments were performed as follows: VTA brain sections were incubated for 1 hour at 4°C in a blocking solution of phosphate-buffered saline (PBS) containing 3% bovine serum albumin (BSA, Sigma; A4503) (vol/vol) and 0.2% Triton X-100 (vol/vol), and then incubated overnight at 4 °C with a mouse anti-tyrosine hydroxylase antibody (anti-TH, Sigma, T1299) at 1:500 dilution, in PBS containing 1.5% BSA and 0.2% Triton X-100. The following day, sections were rinsed with PBS, and then incubated for 3 hours at 22-25 °C with Cy3-conjugated anti-mouse and secondary antibodies (Jackson ImmunoResearch, 715-165-150) at 1:500 in a solution of 1.5% BSA in PBS, respectively. After three rinses in PBS, slices were wet-mounted using Prolong Gold Antifade Reagent (Invitrogen, P36930). Microscopy was carried out with a fluorescent microscope, and images captured using a camera and analyzed with ImageJ. In the case of optogenetic experiments on DAT^iCRE^ mice, identification of the transfected neurons by immunohistofluorescence was performed as described above, with the addition of 1:500 Chicken-anti-GFP primary IgG (ab13970, Abcam) in the solution. A Goat-anti-chicken AlexaFluor 488 (1:500, Life Technologies) was then used as secondary IgG. Neurons labelled for TH in the VTA allowed to confirm their neurochemical phenotype, and those labelled for GFP to confirm the transfection success.

### Intracranial self-stimulation (ICSS) bandit task

#### Behavioral set up

The ICSS bandit task took place in a circular open field with a diameter of 68 cm. Three explicit square-shaped marks (1x1 cm) were placed in the open field, forming an equilateral triangle (side=35 cm). Entry in the circular zones (diameter=6 cm) around each mark was associated with the delivery of a rewarding ICSS stimulation. Experiments were performed using a video camera, connected to a video-tracking system, out of sight of the experimenter. A LabVIEW (National Instruments) application precisely tracked and recorded the animal’s position with a camera (20 frames/s). When a mouse was detected in one of the circular rewarding zones, an electrical stimulator received a TTL signal from the software application and generated a 200 ms-train of 0.5-ms biphasic square waves pulsed at 100 Hz (20 pulses per train). ICSS intensity was adjusted, within a range of 20 to 200 µA, during training (see training contexts) and then kept constant, so that mice would achieve between 50 and 150 visits per session (5min duration) for two successive sessions, and then kept constant for all the experiment. The constant motivational level insured by ICSS alleviated the need for a stimulus to repeat the behavior. Mice with insufficient scores in the PS and DS (<40 visits despite increasing the intensity to a maximum of 200 µA) were excluded.

#### Baseline behavior

Prior to the ICSS bandit task, three control sessions were performed. First, spontaneous neuronal activity was recorded in the mice home-cages for 10 minutes. Second, neuronal activity was recorded while random ICSS were delivered to the mice in its home-cage, to assess the direct effect of the stimulation onto neuronal activity. Third, behavioral and neuronal activity were recorded for 30 min, while the mice were exploring the open-field for the first time (“habituation”, without the presence of the three rewarding locations).

#### Training context

The training consisted of two context s: the deterministic context (D) and the probabilistic context (P), consisting of 10 daily sessions of 5 min for the DS and 10 min for the PS. In the DS, all zones were associated with an ICSS delivery (P=100%). However, two consecutive rewards could not be delivered on the same location, which motivates mice to alternate between locations. In the PS, the zones were associated with three different probabilities (P=25%, P=50%, P=100%) to obtain an ICSS stimulation. The probabilities’ locations were pseudo-randomly assigned per mouse. Animals successively make the task in DS and then in PS.

#### Data acquisition per experimental group

For optogenetics experiments, the DAT^iCRE^ mice (n=16) completed the training, followed by a schedule of 4 days of paired sessions with photo-stimulation (ON) alternated with days without photostimulation (OFF). The averages of the ON and OFF days were compared in a paired manner.

#### Optogenetics experiments

For optogenetic experiments on freely moving mice, an optical fiber (200 µm core, NA=0.39, Thor Labs) coupled to a ferule (1.25 mm) was implanted just above the VTA ipsilateral to the viral injection (coordinates from bregma: AP -3.1 mm, ML ±0.5 mm, DV 4.4 mm), and fixed to the skull with dental cement (SuperBond, Sun Medical). The behavioral task began at least 4 weeks after virus injection to allow the transgene to be expressed in the target dopamine cells. An ultra-high-power LED (520 nm, Prizmatix) coupled to a patch cord (500 µm core, NA=0.5, Prizmatix) was used for optical stimulation (output intensity of 10 mW). Optical stimulation during the behavioral experiment was continuously delivered for 500 ms, starting 100 ms after animal’s detection in a location. The ON and OFF schedule (OFF-ON-OFF-ON-OFF) was following the last week of deterministic training. The optical stimulation cable was plugged onto the ferrule during 5 experimental sessions to prepare the animals and control for latent experimental effects.

#### Intracranial injections of muscimol

A solution of muscimol (TOCRIS) (0.5µg/µL) was infused in the PFC over 20-30 minutes before the beginning of the ICSS bandit task experiment. The bilateral infusion of 0.4µL was performed at a rate of 0.2µL/min using a double injector (Univentor). Before each experiment session, a double injection cannula (2.5 mm, 0.5 mm projection) was inserted into the implanted bilateral cannula guide (length below pedestal 2.5 mm). The injection cannula was connected to a multi-syringe pump (Univentor) that allowed saline or muscimol injection. The saline and muscimol schedule (saline-muscimol-rest-saline-muscimol) was following the last week of probabilistic training. The injection system was plugged onto the cannula guide before 5 experimental sessions to prepare the animals and control for latent experimental effects.

#### Behavioral measures

For all groups of mice, the trajectory was smoothed using a triangular filter allowing the determination of speed profile, which corresponds to instantaneous speed as a function of time, and time of maximal speed within a trial. The following measures were analyzed in the DS and compared in the PS, as well as in the DS for the OFF vs ON Jaws experiment, or in the PS for the Sal vs Mus experiment: i) number of visits, ii) time-to-goal, iii) choice repartition (proportion of visits p_25_, p_50_ and p_100_), iv) percentage of directional changes (n^th^ visit=n^th^ visit+2). Furthermore, the ICSS bandit task can be seen as a Markovian decision process. Every transition between zones can be considered as a binary choice between two probabilities, since the occupied zone cannot be reinforced twice in a row. The sequence of choices per session is summarized by the proportional result of the sum of three specific binary choices (or gambles, i.e., total visits zone 1/total visits zone 1+2). The three gambles (G) were named after the point on which the mouse is positioned at the time of the choice: G_25_=100 % vs 50 %, G_100_=50 % vs 25 % and G_50_=100 % vs 25 %.

Locomotor activity toward the rewarding locations was measured in terms of time-to-goal, dwell time and time to maximal speed. Time-to-goal measures the duration between one location and the next one. The speed profile corresponds to the instantaneous speed as a function of time (20 frames per s). The dwell time is defined as the duration between the end of the 200 ms period (corresponding to the eventual ICSS duration) in the last rewarding location and the moment when the animal’s speed is greater than 10 cm s^−1^. The time to maximal speed is the time at which the speed profile attains its maximal value. We compared general linear regression models (GLM) of the time-to-goal with increasing number of explanatory variables (with Bayesian information criterion). Best explanatory variables were whether the animal performed a U-turn, the dwell time, and the time to maximal speed (minus the dwell time to remove its additive influence). We regularized the GLM for correlated terms using ridge regression, insuring that each predictive variable exerted an uncorrelated effect on the time-to-goal. We finally checked that each parameter had a significant influence (p<0.05) on the time-to-goal for each animal.

#### Modeling

The location choice in these gambles reflects the balance between exploitative (choosing the most valuable option) and exploratory (choosing the least valuable option) choices. With a softmax based decision-making model fitted in the laboratory, we computed three parameters: the value sensitivity or inverse temperature (the power to discriminate between values in a binary choice), the uncertainty bonus (the preference for expected uncertainty, considering the reward variance of every option in a binary choice) and the motor cost to do a directional change (a decrease in the location value if it requires to go back to the previous location). Decision-making models determined the probability *Pi* of choosing the next state i, as a function (the “choice rule”) of a “decision variable”. Because mice could not return to the same rewarding location, they had to choose between the two remaining ones. Accordingly, we modeled decisions between two alternatives labelled A and B and used a softmax choice rule defined by P_A =_1 / (1+e^-ß(vA–vB)^) where β is an inverse temperature parameter reflecting the sensitivity of choice to the difference between decision variables and *Vi* the value of an option. The value *V* of an option is modelled as the expected (average) reward + expected uncertainty + U-turn cost ^16, 30^. This compound value is then nested in the softmax choice rule, given a 6*3 matrix that described the probability of a choice between A, B and C (the three locations) depending on the two previous choices. As an example, in the probability to choose (A, B, C) after performing the sequence BA, the value is given by (0, pb +φpb*(1-pb)-κpc + φ*pc*(1-pc)) while after the sequence CA the value is given by (0,pb+φ*pb*(1-pb),pc+φ*pc*(1-pc) -κ) (same for AB, CB and AC, BC). The free parameters of the model were fitted by maximizing the data likelihood. Given a sequence of choice c=c1..T, data likelihood is the product of their probability (given by Equation 1) ^80^. We derived Bayesian Information Criterion from the likelihood and used it to compare the full model with simpler ones, i.e. a softmax model in which choices only depend on expected value (φ and κ=0) and a softmax model in which choices depend on expected value and motor cost (χπ=0). We also checked that simpler models (null model of random choice, null model with a motor cost, epsilon-greedy with constant exploration) did not provide a better fit. We used the *fmincon* function in Matlab to perform the fits, with the constraints that β ∈]0,10], φ ∈]-1,5] and κ ∈]0,5].

#### Statistical analysis

All statistical analyses were computed using Matlab and Python with custom programs. Results were plotted as a mean ± s.e.m. The total number (n) of observations in each group and the statistics used are indicated in figure legends. Classical comparisons between means were performed using parametric tests (Student’s T-test, or ANOVA for comparing more than two groups) when parameters followed a normal distribution (Shapiro test P>0.05), and non-parametric tests (here, Wilcoxon or Mann-Whitney) when the distribution was skewed. Multiple comparisons were Bonferroni corrected. Probability distributions were compared using the Kolmogorov–Smirnov (KS) test, and proportions were evaluated using a chi-squared test (χ²).

### Electrophysiological recordings

All extracellular potentials recordings were performed using a digital acquisition system (Digital Lynx SX; Neuralynx) together with the Cheetah software. Broadband signals from each wire were filtered between 0.1 and 9000 Hz and recorded continuously at 32 kHz.

#### Multi-unit activity recordings

To extract spike timing, signals were band-pass-filtered between 600 and 6000 Hz and sorted offline. Spike clustering was cross-validated by using both SpikeSort3D (Neuralynx) and custom-written Matlab (The Mathworks) routines. The electrophysiological characteristics of VTA neurons were analyzed in the active cells encountered by systematically moving down the multi-electrodes.

#### Local-field potential recordings

To extract low-frequency variations of extracellular potential, signals were low-pass-filtered below 300 Hz.

#### Population firing

To extract spike timing of the neuronal population, signals were band-pass-filtered between 600 and 6000 Hz and sorted offline. Because population firing originates from bipolar electrodes with only one recording wire, no clustering could be considered.

### Electrophysiological data analysis

#### Identification of DA cells

Extracellular identification of putative DA neurons (pDAn) was based on their location as well as on a set of unique electrophysiological properties that characterize these cells in vivo: 1) a typical triphasic action potential with a marked negative deflection; 2) a characteristic long duration (>2.0 ms) action potential; 3) an action potential width from start to negative trough >1.1 ms; 4) a slow firing rate (<12 Hz) with an irregular single spiking pattern and occasional short, slow bursting activity. Putative GABA neurons were characterized by a characteristic short duration of action potential from start to negative trough (<1.0 ms), and a high firing rate (>12 Hz). D2 receptors (D2R) pharmacology was also used for confirming the DA neurons identification: after a baseline period (5 min) and a saline (10 min) injection, quinpirole (1mg/kg, D2R antagonist) was injected (30 min recording), followed by an eticlopride (D2R agonist) injection (1mg/kg, 10 min recording). Since most DA, but not GABA neurons, express inhibitory D2 auto-receptors, neurons were considered as pDA neurons if quinpirole induced at least 30% decrease in their firing rate, while eticlopride restored firing above the baseline. Nevertheless, as continuous D2 pharmacology could have affected both baseline DA neurons firing and decision-making ^81^, we allowed the mice to recover two days after this experiment. We thus performed pharmacological confirmation (1) when first encountering a putative DA neuron in a given mouse or (2) at the end of the week if at least one putative neuron was present during the behavioral experiment. Neurons were considered as pDAn only if they responded to the pharmacology, or if they presented electrophysiological characteristics defined above and were recorded between two positive pharmacological experiments.

#### Firing analysis

Spontaneous DA cell firing was analyzed with respect to the average firing rate and the percentage of spikes within bursts (%SWB, number of spikes within bursts, divided by total number of spikes). Bursts were identified as discrete events consisting of a sequence of spikes such that: their onset is defined by two consecutive spikes within an interval <80 ms and they terminated with an interval >160 ms. Phasic activity is defined as spikes falling into bursts, while tonic activity comprises spikes outside bursts. Peri-event time histograms (PETH) for normalized activity were constructed based on 1 ms-bins rasters, convolved with a Gaussian kernel (100 ms), divided by the neuron basal firing rate (to compare DA neurons with firing rates from 1 to 10Hz). Normalized PETH were sorted according to the *preceding* event (reward or omission) in Figures 2 and 4, and to the probability of reward associated with the *next* location in Figure 5. Phasic activity from these PETH was defined as the firing rate during a 500ms time window (usually 300-ms-800ms after last location entry unless stated in the Results). We checked that the results did not depend on the exact time window by systematically shifting the beginning (100ms to 500ms) and duration (300ms to 800ms) of the time windows by 50ms bins. Encoding of reward uncertainty by PFC multi-unit activity was also assessed through an enrichment analysis: we determined for which reward probability of target location the PFC population activity was the highest, intermediate and lowest. PFC phasic activity was considered to encode uncertainty if it was highest for 50%, intermediate for 25%, and lowest for 100% probability. The proportion of PFC activity encoding uncertainty was compared to expected proportion (there are 6 possible orders when sorting activities related to 3 events, giving 16.7% as expected proportion).

#### Wavelet analysis

Because extracellular field potentials (EFP) are non-stationary signals, they are transformed offline using a Morlet wavelet transform (center frequency = 0.6 and bandwidth = 1). This process is defined as the convolution product between the EFP signal and dilated forms of wavelets normalized to 1 ^82^. EFP signal was expressed in z-score units in Figure 2. For each channel, the z-score normalization used the mean and the standard deviation from the 2s period preceding the location entry (LE). In Figures 4, 5 and 7, EFP signal was also band-pass filtered in the θ (7-14 Hz) or δ (3-6 Hz) frequency band and normalized for each channel with the mean power in each frequency band. The cross-spectra (cross-correlograms between OFC and PFC power spectra) in Supp. Figure 2 were computed for brain regions of the same hemisphere and per animal. The wavelet coherence (normalized spectral covariance) between the EFP from the OFC and the one from the PFC was computed by smoothing the product of the two wavelet transforms over time (window for time smoothing = 0.2s) and over scale (pseudo-frequency) steps (window for scale smoothing = 2 Hz).

---

**Supplementary Figure 1:**
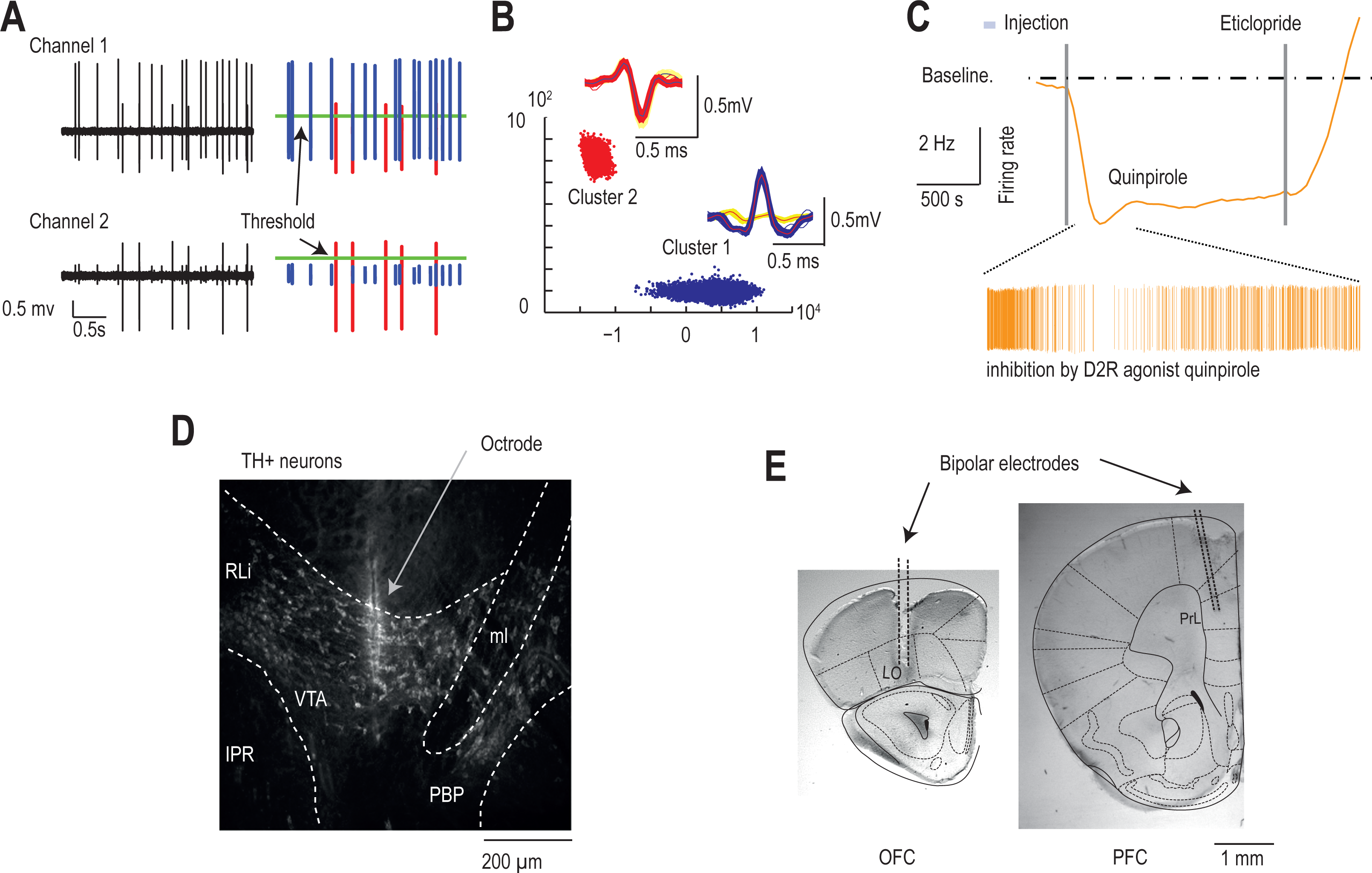
Histological and pharmacological confirmation of dopamine and cortical recordings. **(A)** Band-pass filtered (600-6000Hz) extracellular traces showing spike detection thresholds for an example of a pDAn. **(B)** Illustration of single-unit clustering with principal component representation of spike characteristics with color-coded clusters corresponding to two simultaneously recorded neurons. Insets: spike waveforms (different channels superimposed) from the two clustered neurons. **(C)** Pharmacological confirmation of the dopaminergic nature of recorded neurons. Neurons were considered dopaminergic if inhibited by intra-peritoneal injection of D2R agonist quinpirole followed by reactivation by D2R antagonist eticlopride. **(D)** Histological confirmation of electrode placement with TH staining of dopaminergic cells within the VTA. **(F)** Histological confirmation of electrode placement in orbitofrontal (OFC) and prefrontal (PFC) cortices.

**Supplementary Figure 2:**
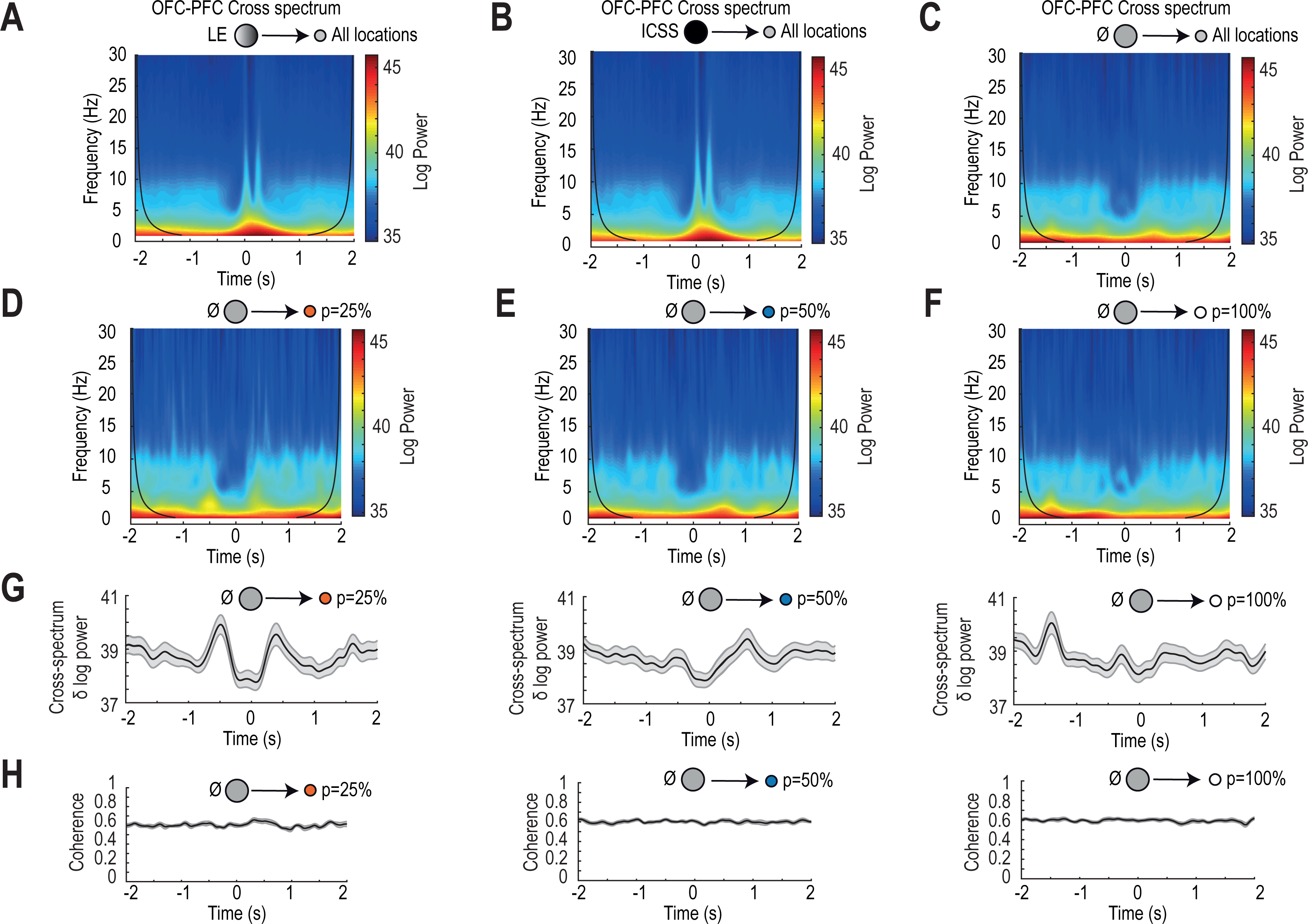
Cross-spectrum and coherence analyses. From A to F: Time and frequency resolved cross-spectrum power density (using complex Morlet wavelet transforms of OFC and PFC EFP) around the reference time (location entry, see below). Reference (0s) time is the time of all location entries **(A)**, ICSS, **(B)** and reward omissions **(C, D, E, F).** Trials are sorted according to the reward probability of the target location: all locations (A, B, C), 25% (D), 50% (E), and 100% (F). **(G)** Time-resolved cross-spectrum power for the 3-6 Hz frequency around reward omissions depending on the reward probability of target location (25% (left), 50% (middle), and 100% (right)). **(H)** Same as (G) for coherence.

**Supplementary Figure 3:**
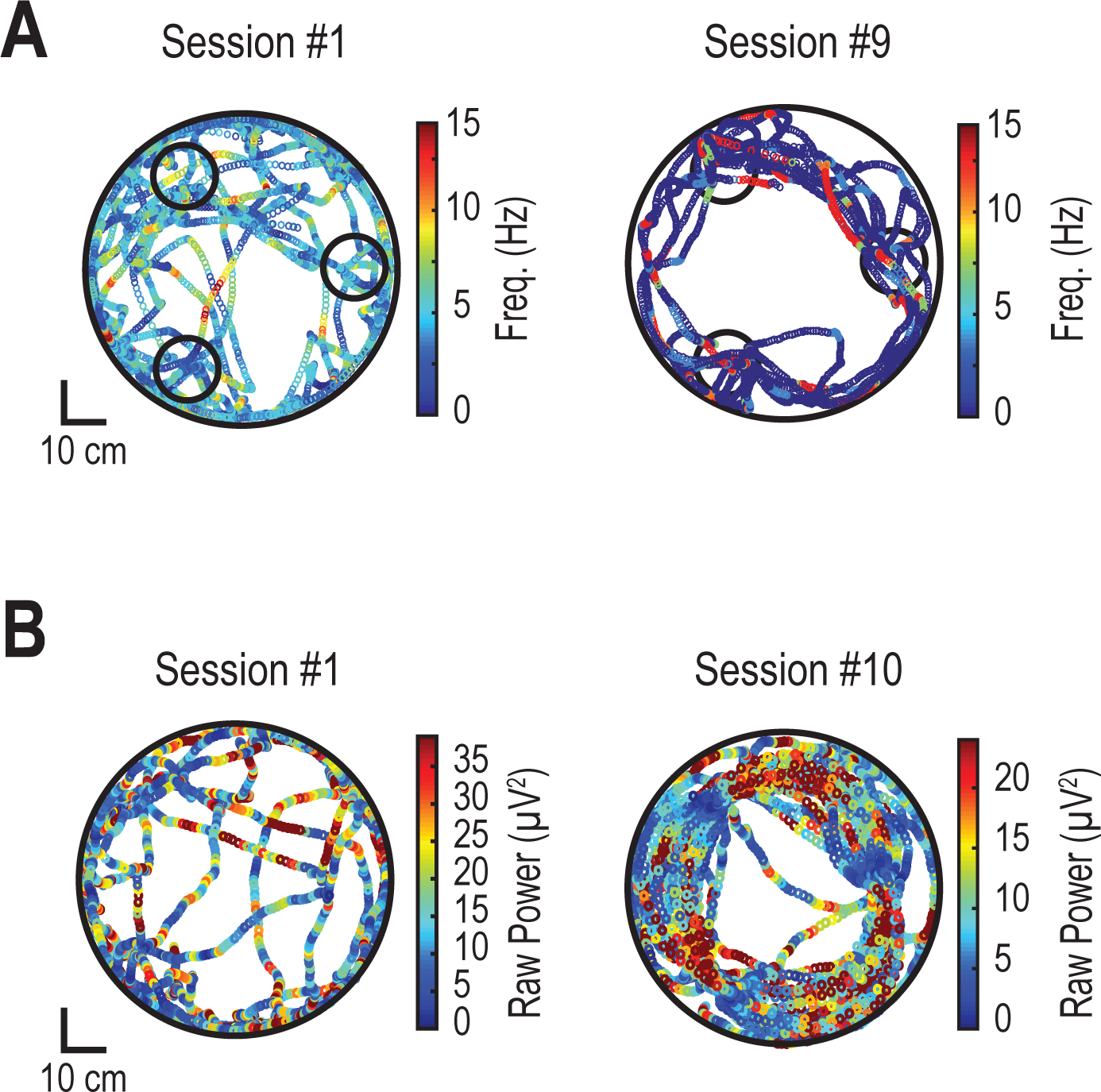
Reorganization of spiking and oscillatory activity during learning. **(A)** Representative examples of animal trajectory with corresponding instantaneous pDAn cell firing frequency (color coded), in the first session (left), and late session (right) of the D context, showing a reorganization of spiking at particular timings together with a shift from tonic to phasic bursting activity. **(B)** Representative examples of animal trajectory with corresponding OFC theta oscillation raw power (color coded), in the first session (left), and late session (right) of the D context, showing an increase in OFC theta oscillations at particular timings.

**Supplementary Figure 4:**
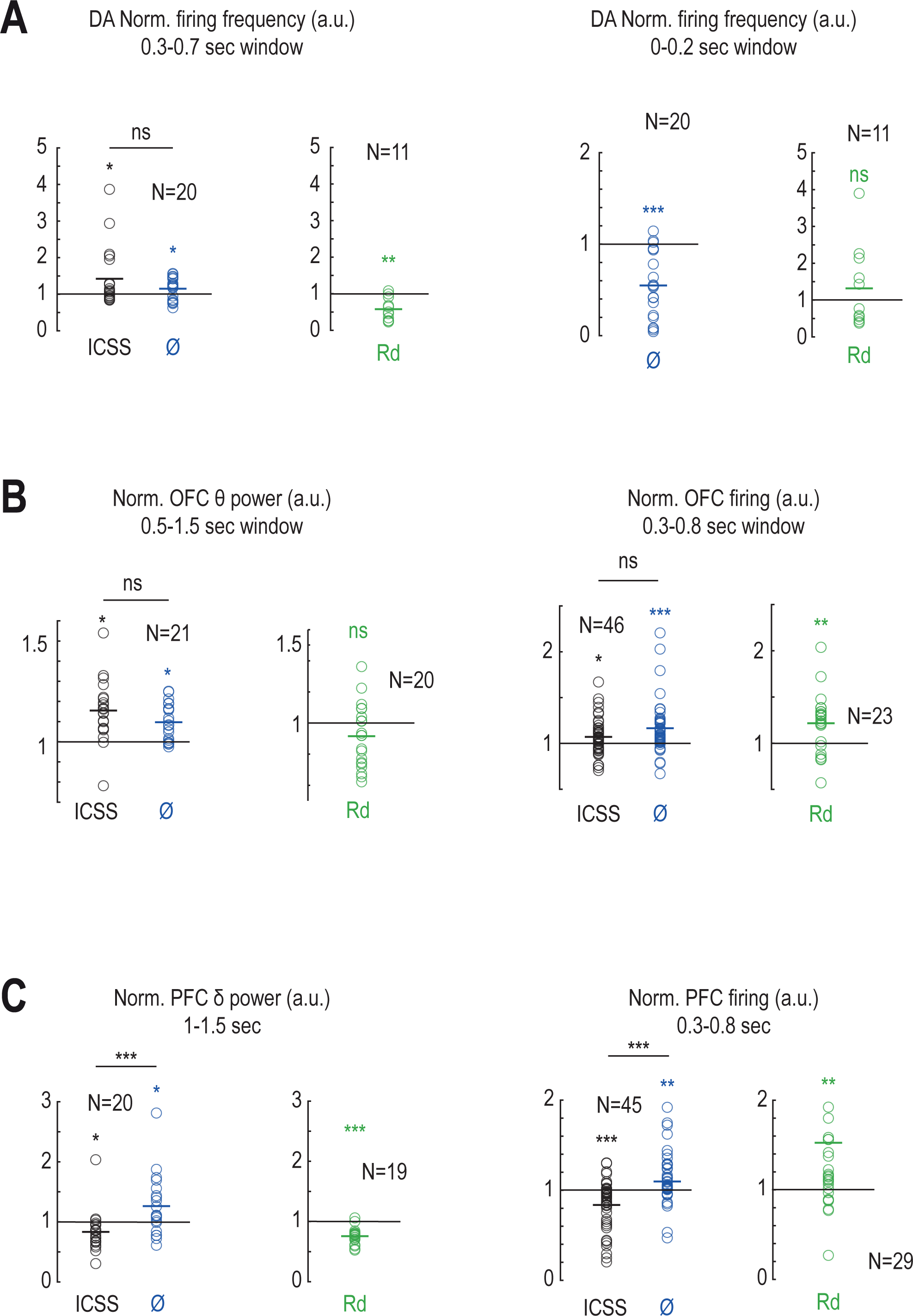
Quantification related to Figure 4. **(A)** Left: Quantification of dopamine firing activity after location entry (over 0.3-0.8s): intra-cranial self-stimulation (black, ICSS), reward omission (blue, ø) and random ICSS in the rest box (green, Rd). Mean pDAn firing after ICSS: two-sided Wilcoxon signed rank test W_(19)_=160 p=0.04; After omission: Student t-test T_(19)_=2.26, p=0.04; After Rd: Student t-test T_(10)_=-4.54 p=0.0011; Difference between ICSS and omission conditions: ns paired Student t-test T_(19)_=1.42. Right: Quantification of dopamine firing activity during the time window of ICSS (over 0-0.2s) for reward omission (blue, ø) and after random ICSS in the rest box (green, Rd). Mean pDAn firing during omission: Student t-test T_(19)_=-5.49, p<0.001. **(B)** Left: Quantification of θ (7-14 Hz) OFC power after location entry (over 0.5-1.5s): intra-cranial self-stimulation (black, ICSS), reward omission (blue, ø) and random ICSS in the rest box (green, Rd). Mean θ power over after ICSS: Student t-test T_(20)_=4.76 p<0.001; After omission: Student t-test T_(20)_=4.86 p<0.001; After Rd: ns Student t-test T_(19)_=-1.86, p=0.08; Difference between ICSS and omission conditions: ns paired ranked Student t-test T_(20)_=-1.51. Right: Quantification of OFC firing activity after location entry (over 0.3-0.8s): intra-cranial self-stimulation (black, ICSS), reward omission (blue, ø) and random ICSS in the rest box (green, Rd). Mean OFC firing after ICSS: Student t-test T_(45)_=2.49, p=0.02; After omission: two-sided Wilcoxon signed rank test W_(45)_=942, p<0.001; After Rd: Student t-test T_(22)_=3.29, p=0.003; Difference between ICSS and omission conditions: ns paired ranked Student t-test T_(45)_=-1.88. **(C)** Left: Quantification of δ (3-6 Hz) PFC power after location entry (over 1-1.5s): intra-cranial self-stimulation (black, ICSS), reward omission (blue, ø) and random ICSS in the rest box (green, Rd). Mean δ power after omission: Student t-test T_(19)_=2.29 p=0.033; After Rd: Student t-test T_(18)_=-7.95, p<0.001; Difference between ICSS and omission conditions: paired two-sided Wilcoxon signed rank test W_(19)_=15, p<0.001, Δ=0.43 a.u. Right: Quantification of PFC firing activity after location entry (over 0.3-0.8s): intra-cranial self-stimulation (black, ICSS), reward omission (blue, ø) and random ICSS in the rest box (green, Rd). Mean PFC firing after omission: two-sided Wilcoxon signed rank W_(44)_=760, p=0.002; After Rd: mean PFC firing over 0.3-0.8s, two-sided Wilcoxon signed rank W_(29)_=363, p=0.002; Difference between ICSS and omission conditions: paired Student t-test T_(43)_=-4.85, p<0.001, Δ=0.28 a.u.

**Supplementary Figure 5:**
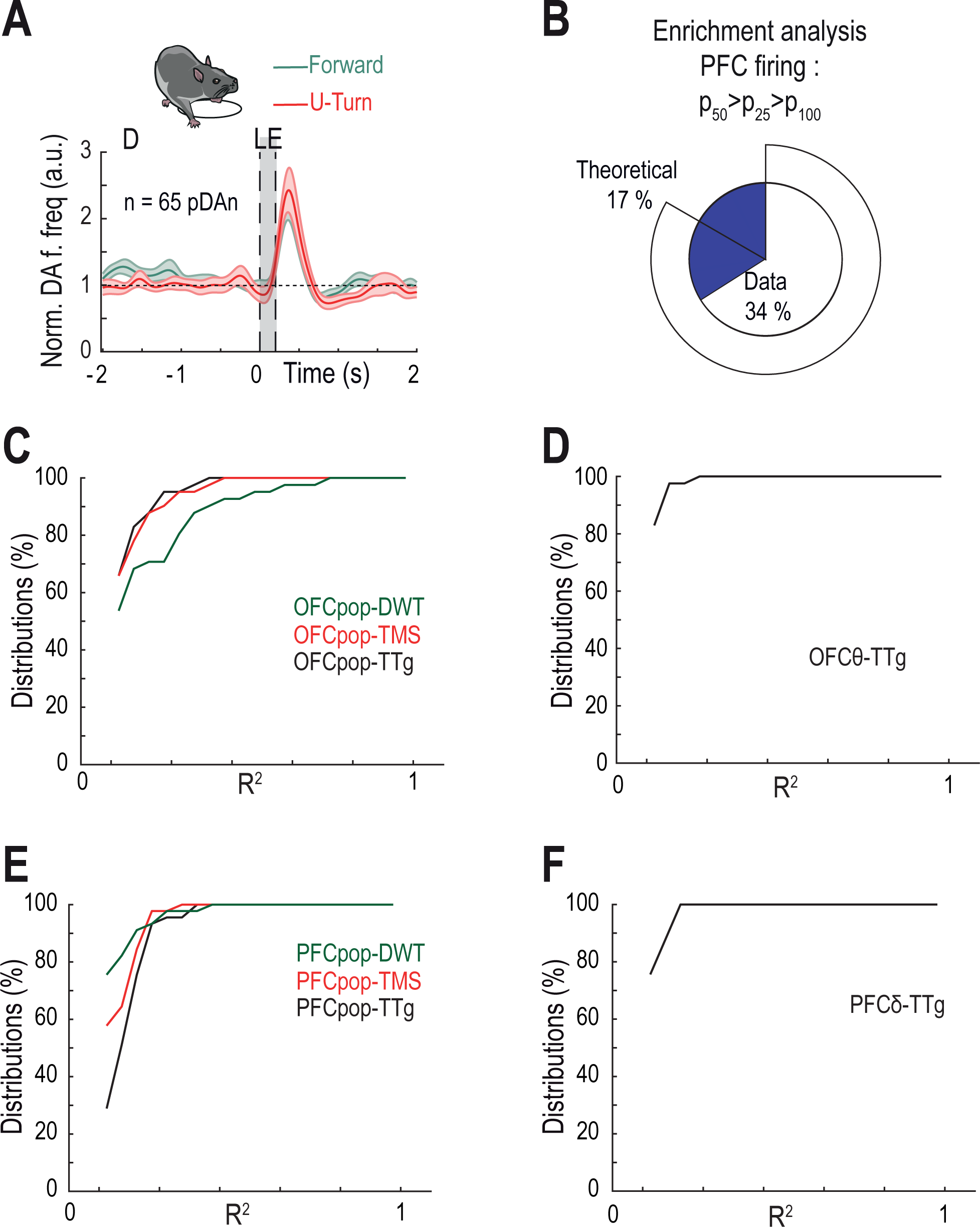
Correlations between neural activity and behavioral timings not shown in Figure 6. **(A)** Normalized firing frequency (a.u.) of pDAn at the end of the D context, centered on location entry (*i.e.* rewards). Trials are sorted according to the direction of the next choice compared to the previous one: U-turn indicating that mice performed a directional change by going back to the previous location (red), in contrast to forward (green). Data are presented as mean ± SEM. **(B)** Proportion of PFC MUA encoding reward uncertainty, *i.e.* displaying a larger activity for p_50_ location than for p_25_, and a larger activity for p_25_ than for p_100_; compared to the theoretical proportion. **(C)** Distribution of correlations coefficients (R^2^) between OFC firing (OFCpop) and time-to-goal (TTG), dwell time (DWT) and time to max speed (TMS). **(D)** Distribution of correlations coefficients (R^2^) between OFC theta oscillation power (OFC θ) and time-to-goal (TTG). **(E)** Distribution of correlations coefficients (R^2^) between PFC firing (PFCpop) and time-to-goal (TTG), dwell time (DWT) and time to max speed (TMS). **(F)** Distribution of correlations coefficients (R^2^) between PFC δ oscillation power (PFC δ) and time-to-goal (TTG).

**Supplementary Figure 6:**
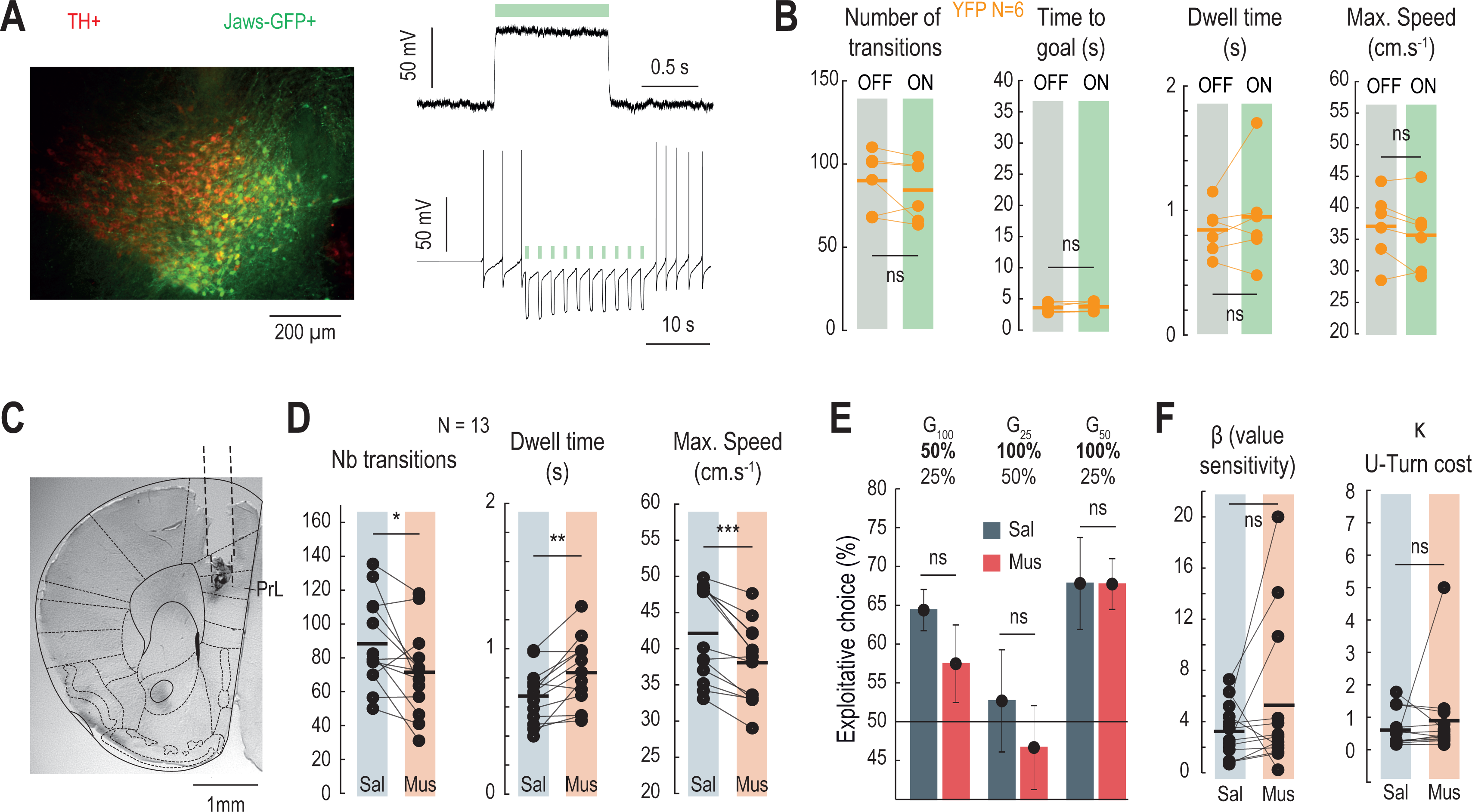
Manipulations of the mesocortical circuit. **(A)** Left: Image of TH+ and Jaws-GFP+ expressing cells in the VTA of a DATi^CRE^ mouse after expression of the inhibitory Jaws opsin. Right: Representative *ex vivo* voltage-clamp (top) and current-clamp (bottom) recordings of a Jaws-expressing VTA DA neuron and illuminated with 1s (top) and 500ms (bottom) continuous light. **(B)** Effect of OFF or ON light stimulations on the number of transitions (*i.e.* of rewards, left) (ns paired two-sided Wilcoxon signed rank test W_(5)_=16), the time to goal (middle left, ns paired Student t-test T_(5)_=-0.91), the dwell time (middle right, ns paired Student t-test T_(5)_=-1.05) and the maximal speed within trials (right, ns paired Student t-test T_(5)_=1.62) in control YFP mice. **(C)** Histological confirmation of intracranial cannula placement in the prefrontal cortex (PFC). **(D)** Effect of muscimol infusions on (left) the number of transitions (*i.e.* of choices), (middle) the dwell time and (right) the maximal speed compared to saline. (Number of transitions: paired Student t-test T_(12)_=2.68, p=0.02, Δ=-16.8; Dwell time: paired Student t-test T_(12)_=-4.01, p=0.0017, Δ=+0.16s; Maximal speed: paired Student t-test T_(12)_=5.28, p<0.001, Δ=-4.06). **(E)** Effect of saline or muscimol infusions on the transition function (the percentage of exploitation) in G_25_, G_100_ and G_50_ gambles. **(F)** Effect of muscimol infusions, compared to saline, on the β parameter (left, ns paired two-sided Wilcoxon signed rank test W_(12)_=19, p=0.068) and the κ parameter (right, ns paired two-sided Wilcoxon signed rank test W_(12)_=41, p=0.79).

